# The exocyst complex is an essential component of the mammalian constitutive secretory pathway

**DOI:** 10.1101/2022.05.26.493223

**Authors:** Conceição Pereira, Danièle Stalder, Georgina Anderson, Amber S. Shun-Shion, Jack Houghton, Robin Antrobus, Michael A. Chapman, Daniel J. Fazakerley, David C. Gershlick

## Abstract

Secreted proteins fulfil a vast array of different functions, including adaptive immunity, cell signalling and extracellular matrix remodelling. In the *trans*-Golgi network, proteins destined for constitutive secretion are sorted into post-Golgi carriers which fuse with the plasma membrane to deliver their contents to the extracellular space. The molecular machinery involved is poorly understood. Here, we have used kinetic trafficking assays and transient CRISPR knock-outs to study the biosynthetic sorting route from the Golgi apparatus to the plasma membrane. Depletion of core-exocyst subunits reduces carrier fusion and causes cargo accumulation in the post-Golgi carriers. Exocyst subunits co-localise with carriers fusing at the plasma membrane and we show that the exocyst complex is recruited directly to these carriers. Abrogation of exocyst followed by kinetic trafficking assays with multiple different soluble cargoes results in cargo accumulation of all tested cargoes. Unbiased secretomics reveals a drastic reduction in the secretion of soluble proteins to the extracellular milieu after knock-out of exocyst subunits. Importantly, the knock-out of exocyst subunits in specialised secretory cell types prevents the constitutive secretion of antibodies in lymphocytes and of the hormones leptin and adiponectin in adipocytes. Together these data identify the exocyst complex as the functional tether of secretory post-Golgi carriers at the plasma membrane and an essential component of the mammalian constitutive secretory pathway.

## Introduction

The complex process of membrane trafficking is fundamental to cellular organisation. Proteins are transported from their site of synthesis in the endoplasmic reticulum to the Golgi apparatus where they are sorted to different subcellular localisations, such as the endolysosomal system or directly to the plasma membrane for secretion^1,2^. In higher eukaryotes, approximately 12% of all proteins are secreted from the cell^3–5^ where they fulfil a vast array of different functions, including cell signalling, the immune response, and extracellular matrix (ECM) remodelling^2^.

Soluble secreted proteins are synthesised in the endoplasmic reticulum. After proper folding, they are trafficked to the Golgi apparatus in COPII carriers where they are glycosylated^1,6–9^. At the *trans*-Golgi apparatus, soluble secreted proteins are sorted into pleomorphic post-Golgi tubular carriers^2^. These carriers are then trafficked directly to the plasma membrane where they fuse and their contents are delivered to the extracellular milieu^2,10^.

The fusion of intracellular carriers is understood to be a two-step process. Molecular tethers either long coil-coiled tethers or multisubunit tethering complexes interact with the carrier prior to the subsequent SNARE-mediated fusion^11,12^. The initial ‘capture’ with the tether is therefore essential for correct vesicle targeting and fidelity of cargo delivery.

Long coil-coiled tethers tend to be large (>60 kDa) and form a coiled-coil domain structure. Examples of long coil-coiled tethering factors include the golgin family of proteins at the Golgi apparatus, and EEA1 on the endosomes^11,13^. They interact with the acceptor compartment on one side and the incoming vesicle on the other ‘bringing’ the vesicle closer to the target membrane^13^.

The second class of membrane tether are multisubunit tethering complexes which include Golgi-associated retrograde protein (GARP)^14^ complex on the *trans*-Golgi network, the conserved oligomeric Golgi (COG)^15^ complex on the *medial*-Golgi, and the homotypic fusion and protein sorting (HOPS)^16^ complex on the lysosomal-endosomal system. These tend to be large multi-subunit assemblies and are sometimes, but not exclusively, complexes associated with tethering containing helical rods (CATCHR) which, when assembled, form helical bundles arranged in tandem through a coiled-coil (CC) region at the N-terminus^17^. Multisubunit tethering complexes have also been found to be important for proper SNARE assembly in addition to vesicle catching^14^.

On the plasma membrane two molecular tethers have been identified. The long coil-coiled protein ELKS (also: ERC, RAB6IP2 or CAST) localises to patches on the plasma membrane termed ‘fusion hotspots’ due to the higher frequency of vesicle fusion events at these sites^18–22^. ELKS was identified as an interactor and probable effector of all three RAB6 isoforms (RAB6A, A’ and B)^19^. ELKS is implicated in secretion of neuropeptide Y in RAB6, MICAL3 and RAB8 positive carriers^23,24^, and synaptic vesicle tethering to the plasma membrane in a neuronal cell model^25^.

The second tether associated with the plasma membrane is the CATCHR protein complex exocyst^26^. Exocyst is an octamer composed of EXOC1-8, and was originally identified in yeast as important for secretion based on its localisation to the plasma membrane and the Sec ^2^. In mammalian cells, exocyst components localise to the Golgi and plasma membrane as well as at vesicle fusion points^27–29^. Although exocyst is essential for endosomal recycling to the plasma membrane^26,29^, the role of exocyst in biosynthetic protein secretion remains unclear. Inhibition of exocyst with antibodies does not affect delivery of *ts*VSV-G, a marker of the secretory pathway, to the plasma membrane^23,27^, however, depletion of *EXOC7* decreases *ts*VSV-G delivery to the plasma membrane^30^. It is therefore not known and remains controversial if exocyst plays a direct role in soluble protein secretion in mammalian cells^2,26^. To investigate the functional machinery in protein secretion, we developed a quantitative trafficking assay to study cell surface delivery from the Golgi apparatus using the retention using selective hooks (RUSH) system. By designing a synthetic type-1 membrane protein based on LAMP1, we can directly observe post-Golgi carriers that co-localise with previously characterised markers and fuse with the plasma membrane. We have used a transient CRISPR-KO system to determine that the exocyst complex is essential for the arrival of these carriers to the plasma membrane. We observe exocyst subunits localising to the post-Golgi carriers on fusion ‘hot-spots’ on the plasma membrane. Kinetic trafficking assays on a set of soluble secreted proteins reveals a broad dependence on exocyst for protein secretion. We performed unbiased proteomics in an endogenous context to exocyst-KO cells and have demonstrated that the exocyst complex is responsible for the majority if not all soluble protein secretion. In addition, we show that important specialised secretory cells require exocyst for the efficient secretion of both antibodies and hormones. We, therefore, define exocyst as the molecular tether for constitutive protein secretion of soluble proteins and as an essential component of the mammalian secretory pathway.

## Results

### Generation of quantitative kinetic cell-surface trafficking assay

In order to study post-Golgi carriers, we have generated a synthetic protein that allows monitoring of protein delivery to the plasma membrane with proper spatiotemporal kinetics. The single-pass type-1 integral membrane protein LAMP1 is localised to the lysosome at the steady-state level. After synthesis in the ER, LAMP1 traffics via the Golgi apparatus to the plasma membrane where it is endocytosed, in clathrin-coated vesicles, to be delivered to the endolysosomal system and finally to the lysosome^1^. Mutations in, or deletion of, the endocytic trafficking motif causes LAMP1 to accumulate on the plasma membrane after exit from the Golgi apparatus in post-Golgi tubular carriers^1^. To monitor the kinetics of trafficking we used the RUSH system^31^. LAMP1ΔYQTI was genetically fused to a streptavidin-binding peptide (SBP) and a fluorescent protein (GFP) and coexpressed with streptavidin fused to the ER-retrieval signal KDEL^32^ in a stable cell line (Fig.1A).

**Figure 1.**
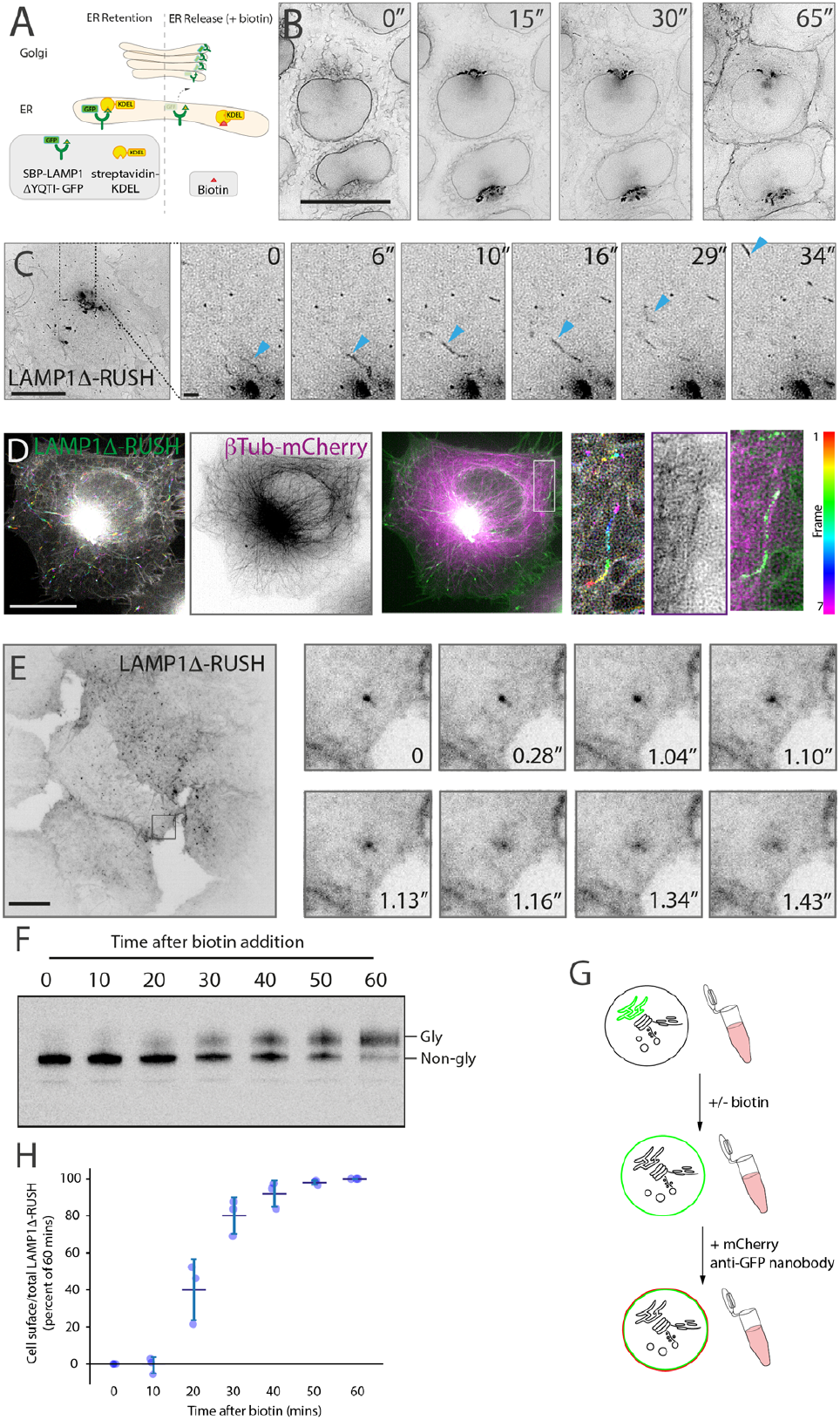
The biosynthetic LAMP1Δ-RUSH (“retention using selective hooks”) reporter system. (**A**) Schematic representation of the RUSH system. By co-expressing the endoplasmic reticulum (ER) hook streptavidin-KDEL with a reporter fused to streptavidin binding peptide-GFP the reporter can accumulate in the ER through the interaction between streptavidin and streptavidin binding peptide. Addition of biotin allows for release of the reporter, which then traffics en masse through the secretory pathway. (**B**) Kinetic analysis of current RUSH cell line based on previous work. The type-1 membrane spanning RUSH reporter LAMP1 Δ-GFP is used to monitor transport through the secretory system. Upon the addition of biotin, LAMP1Δ-RUSH traffics from the ER (0’), to the Golgi apparatus (15’) and then directly to the plasma membrane (30’-45’). Scale bar: 10 *μ*m. (**C**) Lattice-SIM imaging allows the observation of LAMP1 Δ-RUSH leaving the Golgi apparatus in tubular carriers 35 min after biotin addition. In the example image (single micrograph from the time series), the cytosol can be seen full of these tubular structures. Scale bar: 10 *μ*m. (**D**) Time colour coded max projection of LAMP1 Δ-RUSH carriers (represented by the RGB colour bar 0s-6s) shows their trajectory over time along the microtubular network (shown as a max projection). The insert shows a close-up example. Scale bar: 10 *μ*m. (**E**) Plasma membrane TIRF plane showing LAMP1Δ-RUSH post-Golgi tubule fusion. Tubules can be seen as bright spots as they approach and fuse, after which the cargo laterally diffuses on the plasma membrane. Scale bar: 10 *μ*m. (**F**) GFP immunoblot showing increasing LAMP1 Δ-RUSH glycosylation over time after biotin addition. Note that LAMP1 Δ-RUSH starts showing glycosylation from 20 min onwards. Gly = glycosylated LAMP1Δ-RUSH, Non-gly = non-glycosylated LAMP1Δ-RUSH. (**G**) Schematic representation of the RUSH plus cell surface staining protocol developed for FACS analysis. (**H**) Quantitative cell surface assay showing time course arrival of LAMP1 Δ-RUSH to the plasma membrane after biotin addition.

After biotin addition, the LAMP1 ΔYQTI-RUSH (referred to from here as LAMP1 Δ-RUSH) can be observed trafficking with appropriate kinetics (Fig. 1B) as previously observed^1^. Using lattice-SIM live-cell imaging, carriers can be observed budding from the Golgi apparatus (Fig. 1C), trafficking along microtubules (Fig.1D), and using TIRF microscopy we can observe them fuse with the plasma membrane (Fig. 1E). The LAMP1Δ-RUSH is progressively glycosylated after the addition of biotin (Fig.1F). To quantitatively study this secretory route we have developed a FACS-based quantitative cell-surface protein delivery assay (Fig.1G). By using a purified anti-GFP nanobody fused to a fluorescent mCherry^33^ we can quantify delivery to the plasma membrane of the integral membrane protein cargo (Fig. 1H). In summary, the use of the RUSH system with a LAMP1ΔYQTI cargo allows for observation of post-Golgi tubular carriers with appropriate trafficking kinetics and a quantitative read-out.

### RAB6A, ARHGEF10 and RAB8A associate with post-Golgi tubular carriers

To validate that the post-Golgi carriers observed using LAMP1 Δ-RUSH are representative of secretory carriers, we tested if known-markers of these carriers co-localise with LAMP1Δ-RUSH. The small G protein RAB6A/A’ has been observed associated with secretory vesicles that fuse directly with the plasma membrane and has an important role in the fission of the vesicles at the Golgi apparatus and their transport towards the cell surface^23,34^. Lattice-SIM live-cell imaging showed that overexpressed HALO-RAB6A is associated with tubular carriers emerging from the Golgi apparatus, detaching and travelling towards the plasma membrane (Fig.2A). Further, overexpressed HALO-RAB6A colocalised with LAMP1Δ-RUSH at the Golgi apparatus and on these carriers (Fig. 2B). Another small G protein, RAB8A, associates with exocytotic vesicles in a Rab6-dependent manner through the recruitment of the exchange factor ARHGEF1 0^24,35^. Live-cell imaging demonstrated that both HALO-ARHGEF10 and HALO-RAB8A colocalise with LAMP1 Δ-RUSH (Fig. 2C,D), recapitulating previous evidence of a Rab cascade. In conclusion, the post-Golgi carriers observable using LAMP1 Δ-RUSH colocalise with a number of markers for secretory carriers.

**Figure 2.**
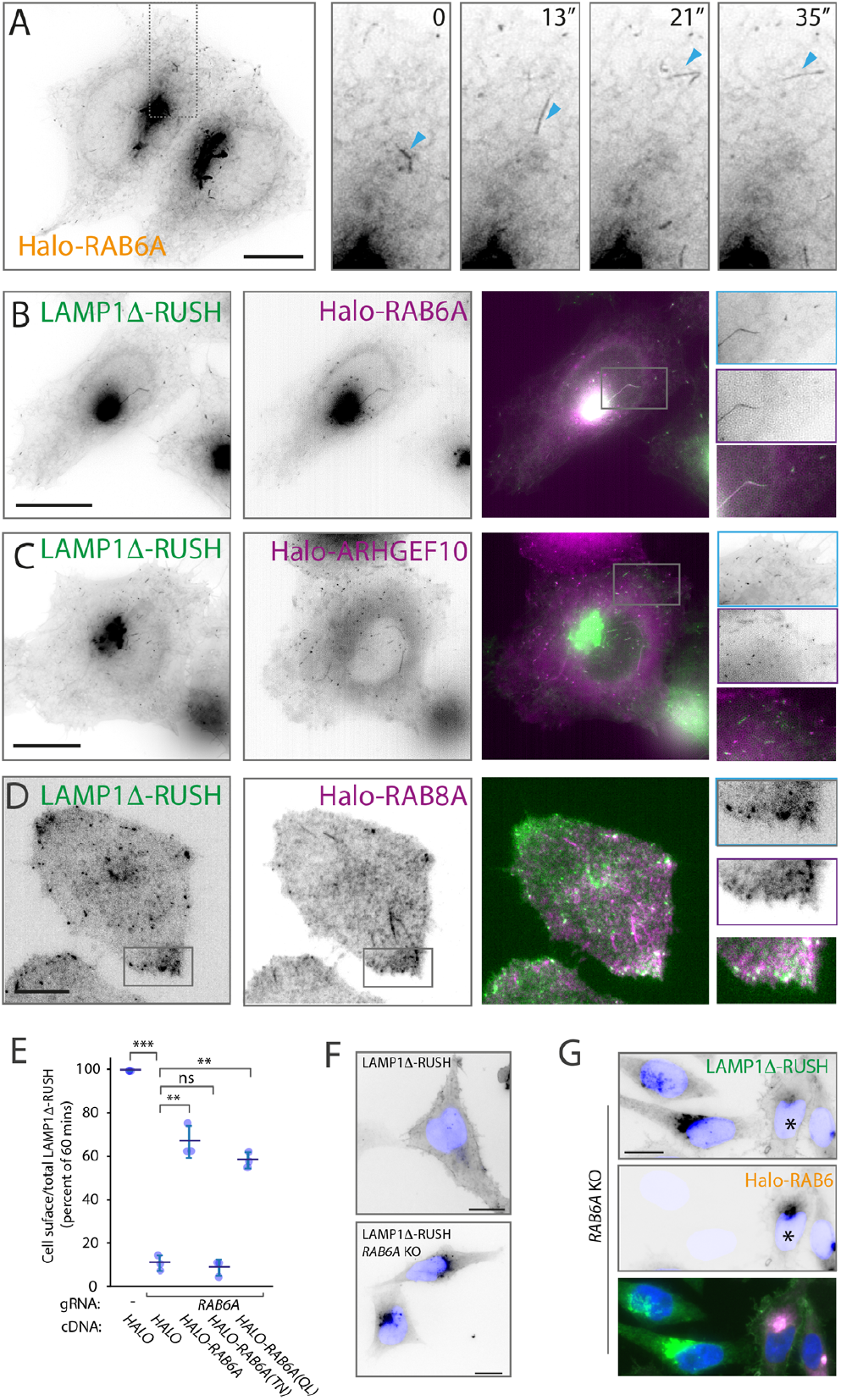
RAB6A, ARHGEF10 and RAB8A colocalise with LAMP1Δ-RUSH post-Golgi carriers. (**A**) Heterologous expression of *HALO-RAB6A* in WT HeLa cells shows RAB6A present at the Golgi and in tubular structures that bud off and travel towards the plasma membrane (single micrograph from a Lattice-SIM time series). (**B**) Lattice-SIM imaging showing HALO-RAB6A colocalising with LAMP1Δ-RUSH tubules (blue) leaving the Golgi and moving towards the plasma membrane (35 min after biotin addition). Scale bar: 10 *μ*m. (**C**) Lattice-SIM imaging showing colocalsation of HALO-ARHGEF10 with LAMP1Δ-RUSH carriers (blue) travelling towards the plasma membrane (35 min after biotin addition). Scale bar: 10 *μ*m. (**D**) Plasma membrane TIRF plane showing HALO-RAB8A colocalising with LAMP1 Δ-RUSH carriers (blue) near their fusion site at the plasma membrane. Scale bar: 10 *μ*m. (**E**) Transient RAB6A-KO dramatically reduces LAMP1 Δ-RUSH at the plasma membrane in a quantitative cell surface assay carried out 35 min post-biotin addition. Note that expression of *HALO-RAB6A WT* (rescue), and constitutively active *RAB6A (QL),* is able to restore plasma membrane expression but not the constitutively inactive form (TN). (**F**) Widefield imaging of LAMP1Δ-RUSH WT and *RAB6A* transient KO, 1 hour after biotin addition. Image shows the LAMP1Δ reporter unable to leave the Golgi in the *RAB6A* KO cells. Scale bar: 10 *μ*m. (**G**) Widefield imaging of LAMP1 Δ-RUSH reporter in *Rab6A*-KO cells expressing *HALO-RAB6A* (*rescue). Upon *RAB6A* heterologous expression, the reporter is at the plasma membrane 1 hour after biotin addition. Scale bar: 10 *μ*m. *P≤0.05; **P≤0.01; ***P≤0.001.

To demonstrate a functional relationship between RAB6 and LAMP1 Δ-RUSH carriers we transiently abrogated RAB6A using CRISPR-Cas9. Loss of RAB6A drastically decreased LAMP1Δ-RUSH trafficking to the plasma membrane (Fig. 2E). Unlike the inactive form of RAB6A (HALO-RAB6A(TN)), overexpression of Rab6A wild-type (HALO-RAB6A) as well as the dominant active form (HALO-RAB6A(QL)) rescued the defect in cell surface delivery of LAMP1Δ-RUSH (Fig. 2E). *RAB6A*-KO caused accumulation of LAMP1 Δ-RUSH in the Golgi apparatus, which was rescued by overexpressing HALO-Rab6A (Fig. 2F-G), showing that Rab6A is required for the generation of LAMP1Δ-RUSH carriers. Together, these results show that markers of secretory carriers colocalise with LAMP1 Δ-RUSH carriers and are necessary for their trafficking from the Golgi, thus demonstrating the LAMP1 Δ-RUSH carriers represent the secretory pathway to the plasma membrane, providing a tractable experimental system to study the secretory pathway.

### Exocyst components associate with fusion hot-spots and are essential for carrier delivery to the plasma membrane

Having established a quantitative system for studying the secretory pathway, we utilised it to study the role of plasma membrane tethers in secretory carrier fusion to the cell surface. The long-coil coiled membrane tether ELKS has been previously associated with the fusion of carriers at the plasma membrane^22^. Consistent with these studies, ELKS colocalises on post-Golgi carrier fusion hot-spots by TIRF microscopy (Fig. S1A). Transient or stable KO of *ELKS* or double KO of *ELKS* and its characterised cofactors or homologs by CRISPR-Cas9 was performed and validated by immunoblot or qPCR (Fig. S2). No significant decrease in cell surface delivery of LAMP1Δ-RUSH to the cell surface could be detected (Fig. S1B-E).

Based on previous evidence in yeast model systems we hypothesised that the octameric protein complex exocyst is essential for the fusion of post-Golgi carriers with the plasma membrane^36–38^. Heterologous expression of exocyst subunit *EXOC1* in the LAMP1Δ-RUSH cell line followed by TIRF microscopy demonstrated the presence of exocyst on ‘hot-spots’ at the plasma membrane co-localising with fusion points of post-Golgi tubules (Fig. 3A). The exocyst is considered an essential component of eukaryotic cells^39,40^, so to avoid lethality and clonal selection artefacts we performed transient CRISPR-Cas9 KO of exocyst components. Using this approach CRISPR-Cas9 abrogation of exocyst components *EXOC3* and *EXOC5* significantly decreased cell surface delivery of LAMP1Δ-RUSH by greater than 60% (Fig. 3B,C). Rescue with overexpression of *EXO-HALO* fusions demonstrated that the defect in cell surface delivery of LAMP1 Δ-RUSH is not due to off-target effects or generalised cell lethality (Fig. 3B,C). Imaging *exocyst*-KO cells demonstrated an accumulation of post-Golgi carriers at the cell tips (Fig. 3D). Interestingly, double knock-out of *ELKS* and *EXOC1* resulted in a significant increase in the phenotype (Fig. S1F), highlighting that ELKS contributes to this pathway. Together these data demonstrate that the exocyst complex is of fundamental importance for cell surface delivery of LAMP1Δ-RUSH carriers.

**Figure 3.**
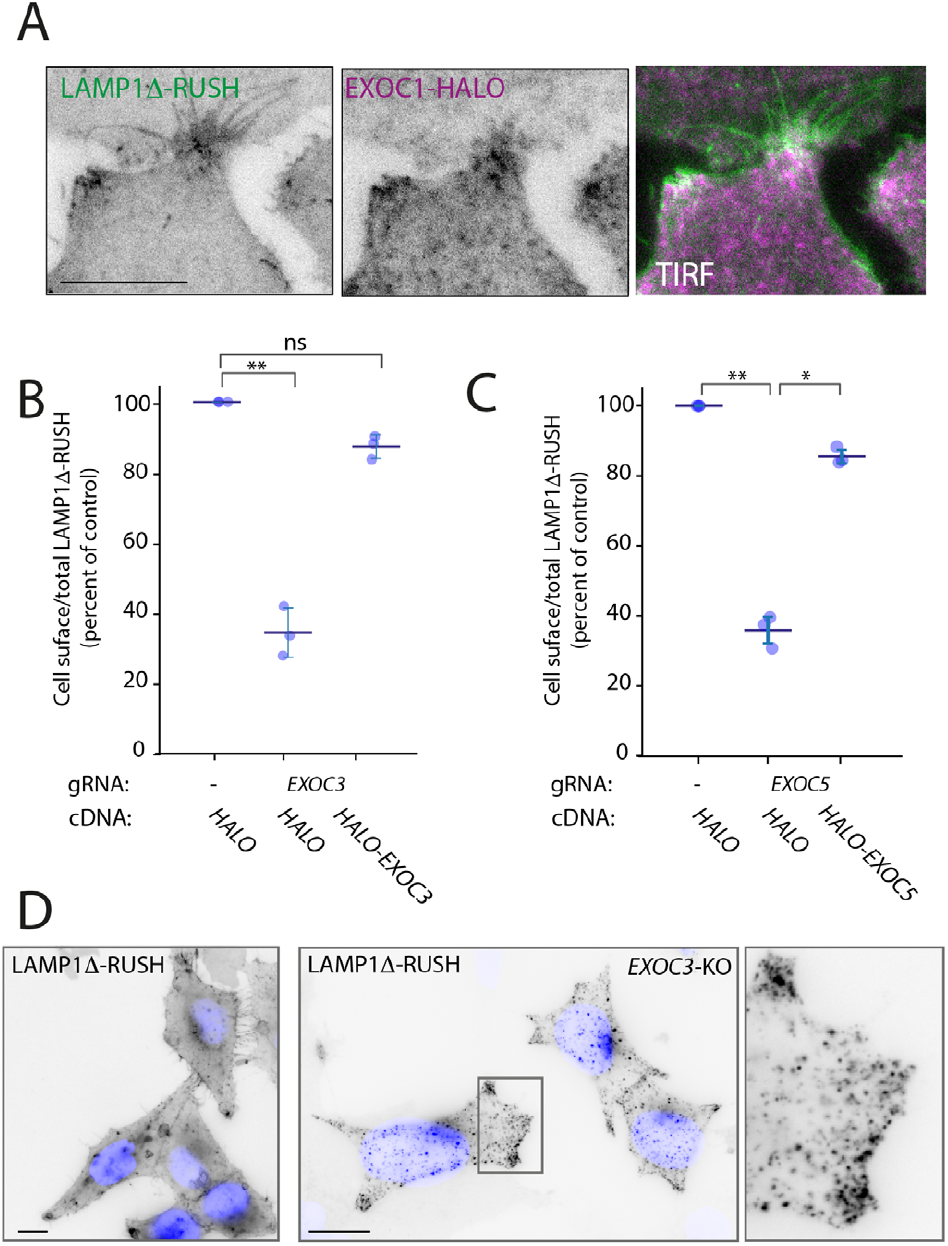
Exocyst components localise to LAMP1Δ-RUSH fusion hot-spots and are essential to plasma membrane delivery. (**A**) TIRF imaging of LAMP1Δ-RUSH (grey) cells expressing heterologous EXOC1-HALO (red) after 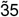 min in biotin. Arrowheads denote fusion sites. Scale bar: 10 *μ*m. (**B**) Cell surface ratio quantification (FACS) showing reduced amounts of LAMP1Δ-RUSH at the plasma membrane after transient *EXOC3* KO and 35 min of biotin exposure. Heterologous expression of EXOC3-HALO recovers this phenotype. (**C**) Cell surface ratio quantification (FACS) showing comparable phenotype and recovery behaviour for EXOC5. (**D**) Widefield imaging of LAMP1Δ-RUSH reporter in *EXOC3* KO cells 1 hour after biotin addition. When compared to WT, EXOC3 KO cells show substantial accumulation of LAMP1Δ-RUSH carriers near the plasma membrane. Scale bar: 10 *μ*m. *P≤0.05; **P≤0.01.

### Exocyst is directly recruited to post-Golgi carriers and multiple exocyst-associated proteins are essential for carrier delivery

To ensure that the effect of *exocyst*-KO on post-Golgi carrier fusion is direct, we tested localisation of exocyst components. By tagging EXOC6 with a HaloTag and expressing it at low levels in our LAMP1Δ-RUSH cell line, we observed EXOC6 at fusion punctae co-localising with LAMP1Δ-RUSH (Fig. 4A). To test the localisation of the other exocyst subunits to the carriers we developed a biochemical approach to localise proteins of interest to post-Golgi carriers. We generated a second LAMP1Δ-RUSH system with the GFP on the cytosolic side of the membrane (Fig. 4B). This fusion had comparable kinetics to the lumenal/extracellular tagged variant. 35 minutes after the addition of biotin there is an accumulation of post-Golgi carriers in the cytosol. We mechanically lysed the cells and immuno-isolated the carriers using GFPtrap beads, an approach we term ‘carrierIP’. Expression of the control HALO resulted in no enrichment after carrierIP, and all core exocyst components tested (EXOC1, EXOC3, EXOC5, EXOC6 and EXOC7) were enriched on post-Golgi carriers (Fig. 4C). These data demonstrate that exocyst is directly recruited to the post-Golgi carriers.

**Figure 4.**
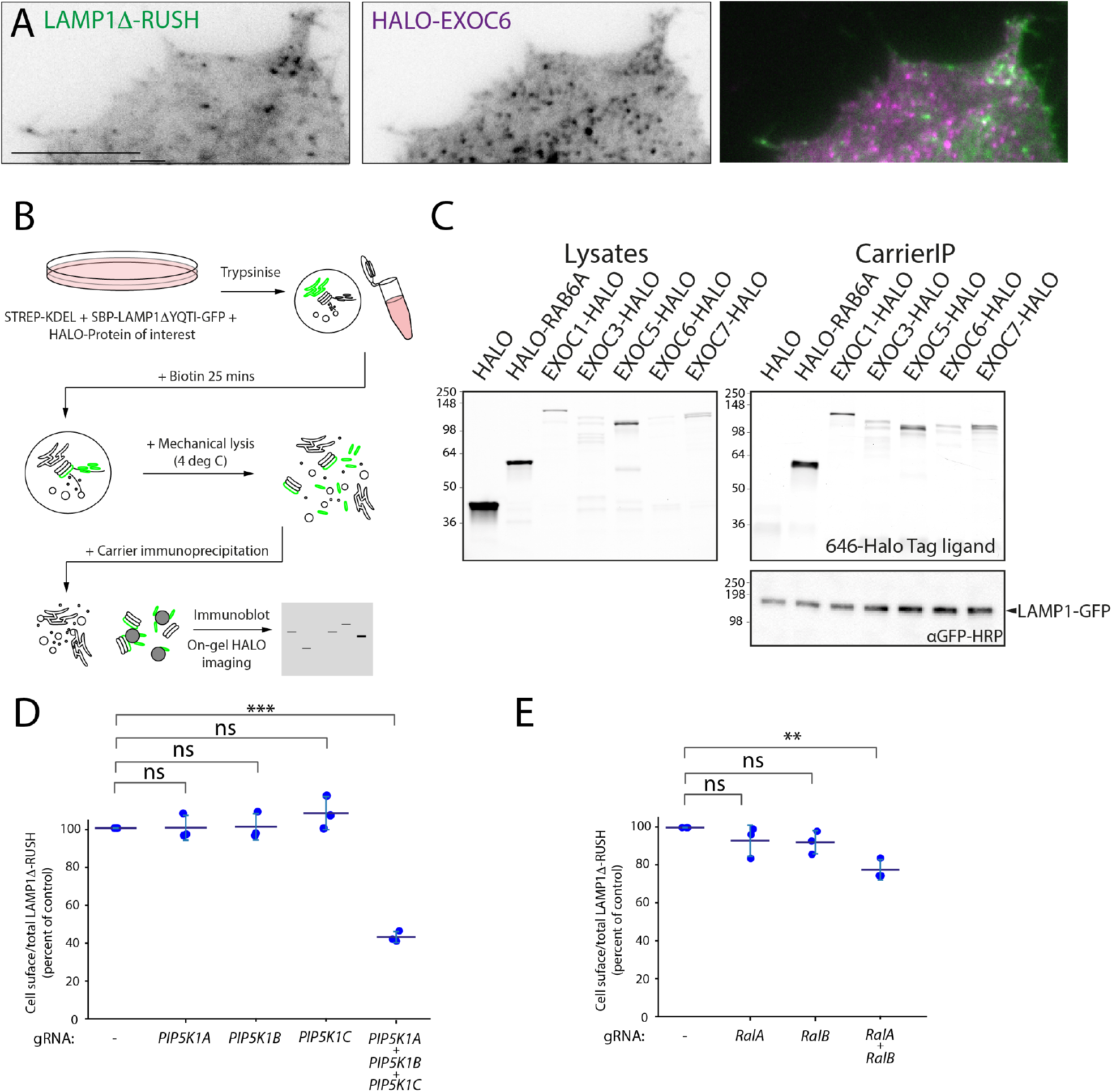
Exocyst is recruited to post-Golgi carriers and multiple associated proteins are essential. (**A**) TIRF imaging showing heterologous expression of HALO-EXOC6 in the LAMP1Δ-RUSH reporter cells. EXOC6 mostly localises to active fusion sites and specifically colocalises with LAMP1Δ-RUSH carriers near the plasma membrane. Scale bar: 5 *μ*m. (**B**) Schematic representation of RUSH carrierIP assay. (**C**) Gels containing resolved proteins from carrierIP assay denoting enrichment of exocyst subunits (HALO-tagged EXOC1, EXOC3, EXOC5, EXOC6 and EXOC7) in LAMP1Δ-RUSH post-Golgi carriers (LAMP1 immunoblot). (**D**) Cell surface ratio quantification (FACS) showing reduced amounts of LAMP1Δ-RUSH at the plasma membrane after transient KO of *PIP5K* homologs and 35 min of biotin exposure. (**E**) Cell surface ratio quantification (FACS) showing amounts of LAMP1Δ-RUSH at the plasma membrane after transient KO of *RALA* and/or *RALB*; and 35 min of biotin exposure. **P≤0.01; ***P≤0.001.

Biochemical studies have identified that the phosphatidylinositol-5 kinase (PIP5K) is important for the recruitment of exocyst to the membrane by modifying phosphatidylinositol to phosphatidylinositol-5 phosphate^41^. There are three PIP5K1 homologues in mammalian cells, PIP5K1A, PIP5K1B and PIP5K1C. Knock-out of either isoform individually had no detectable effect on cell surface delivery of LAMP1Δ-RUSH (Fig. 4D). Triple-transient knock-out of all three isoforms, however, lead to a decrease of around 60% (Fig. 4D), comparable to the loss of exocyst (Fig. 3B-D). The small GTPase Ral has been shown to be another important factor for exocyst recruitment^41^. There are two Rals in mammalian cells, RALA and RALB. Loss of individual Rals had no significant effect on cell surface delivery of LAMP1Δ-RUSH to the cell surface (Fig. 4E). Transient CRISPR knock-out of both subunits significantly decreased cell surface delivery of LAMP1Δ-RUSH. Together these data directly tie the machinery known to be important for exocyst recruitment to post-Golgi carriers to the plasma membrane.

### Exocyst is essential for the secretion of multiple cargoes

There are multiple soluble constitutive cargoes associated with post-Golgi carriers. It is not known if these cargoes undertake different trafficking routes to the plasma membrane. To find which of these potential routes are dependent on exocyst we developed a novel RUSH construct in a PiggyBac transposon system backbone (Fig. 5A) into which we subcloned different soluble secreted cargoes. These cargoes included PAUF (CARTS^42^), CAB45 (sphingomyelin carriers^43^), Collagen X (COLX, RAB6 positive carriers^22^), NUCB1 (a soluble secreted protein) and signal peptide-HALO (synthetic soluble cargo). We generated stable cell lines from these vectors which resulted in all cargoes being retained in the ER at steady state (Fig. S3) and subsequently performed transient CRISPR-Cas9 KO of exocyst component *EXOC3,* as before (Fig. 5B). Widefield imaging demonstrated that all cargoes were effectively held in the ER (Fig. S4A). We then performed RUSH assays and used an unbiased FACS assay to quantitatively monitor the loss of these soluble cargoes. All cargoes demonstrated a loss after RUSH, which was significantly decreased due to the deletion of exocyst (Fig. 5C). Live cell lattice-SIM imaging demonstrated that in *EXOC3*-KO cells, in all conditions, cargoes can be observed accumulating in the cell tips, a phenocopy of the LAMP1 ΔYQTI phenotype (Fig. 5D). Exocyst is therefore essential for the cell surface delivery of all tested soluble, secreted plasma membrane cargoes.

**Figure 5.**
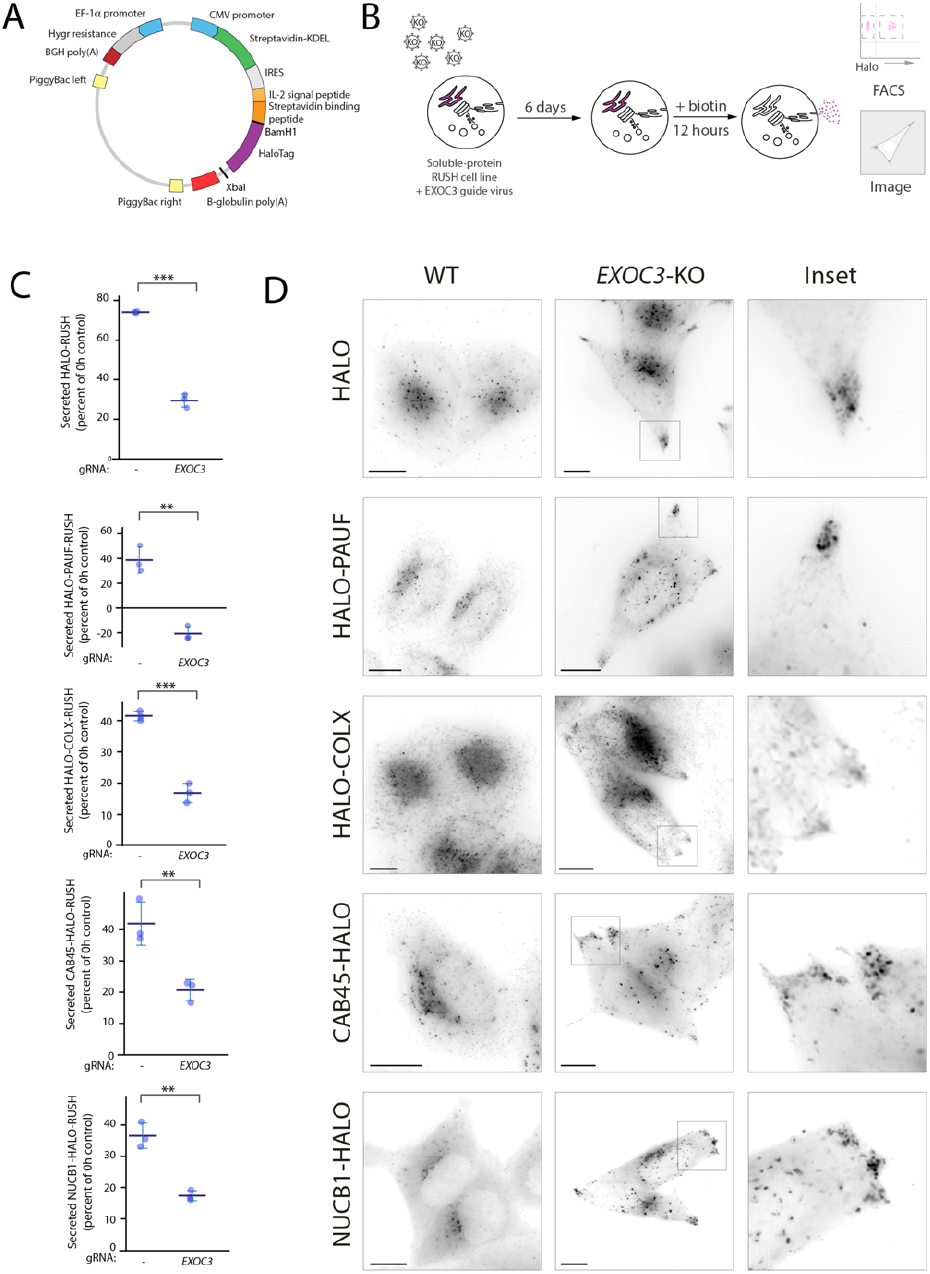
Exocyst is essential for the secretion of various soluble cargoes. (**A**) Schematic of novel PiggyBac transposon RUSH backbone. Genes encoding proteins of interest were either cloned upstream (BamHI) or downstream (XbaI) of HALO. (**B**) Schematic representation of guide infection followed by RUSH and FACS analysis/imaging of reporter protein of interest. (**C**) Percentage of HALO-tagged RUSH proteins secreted over 12 hours. Transient *EXOC3* KO significantly reduces the amount of HALO secreted over this time period. (**D**) Live lattice-SIM imaging of RUSH cargo proteins in *EXOC3* KO cells 12 hours after biotin addition. When compared to WT, EXOC3 KO cells show substantial accumulation of carriers containing cargo proteins of interest near the plasma membrane. Scale bar: 10 *μ*m. *P≤0.05; **P≤0.01; ***P≤0.001.

### Exocyst is essential for secretion of a broad array of soluble secreted proteins

To this point, all experiments have been performed in cells overexpressing proteins in the RUSH system. Thus, the cells are not only overexpressing test cargoes but, in addition, there is a wave of cargo sorting through the cell. To test if endogenous soluble protein secretion is affected in the absence of exocyst, we performed secretomics on HeLa cells after transient CRISPR-Cas9 KO of *EXOC3*. After filtering the datasets bioinformatically for secreted proteins (i.e. containing a signal peptide, no transmembrane domain and no GPI anchor), we see a drastic decrease in the secretory profile of cells depleted of exocyst (Fig. 6A). Fifty-one proteins are significantly (p<0.05) secreted in the WT cells as compared to the KO cells (Table S2). This included all previously tested proteins that are expressed in HeLa cells^44^ (NUCB1, Cab45, and various members of the collagen family). To ensure that this was not due to expression differences in the WT cells compared to the KO cells, we selected several of these (TIMP2, CST3 and PSAP) with commercially available antibodies, to validate these results. As expected, there was an accumulation of these soluble endogenous proteins in *EXOC3-KO* cells. Thus, exocyst is widely essential for the soluble protein constitutive secretory pathway in HeLa cells.

**Figure 6.**
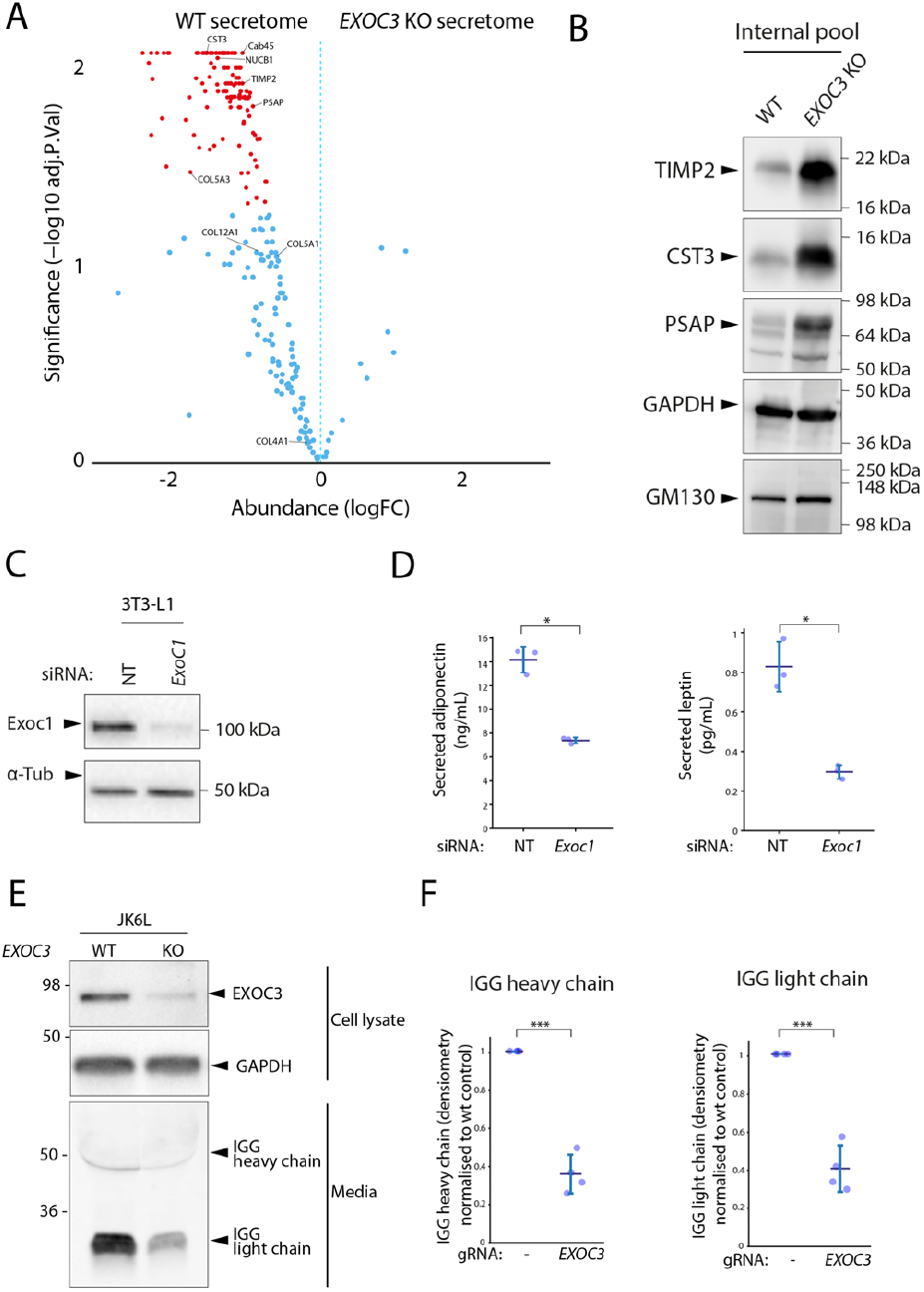
Exocyst is essential for secretion of a broad array of endogenous soluble secreted proteins. (**A**) Volcano plot from the mass spectrometry data on soluble proteins secreted over a period of 6 h by WT HeLa cells, demonstrates that *EXOC3* KO causes a dramatic and significant decrease of protein secretion in these cells. Significantly different proteins are shown in red (p leq 0.01). (**B**) Immunoblot showing intracellular accumulation of selected proteins with downregulated secretion in *EXOC3* KO cells. (**C**) Immunoblot confirming downregulation of EXOC1 in a siRNA-treated population of mouse 3T3-L1 adipocytes. (**D**) Quantification of secreted adiponectin from 3T3-L1 adipocytes siRNA treated with non-targeting (NT) or *Exoc1* 21 h. (**E**) Quantification of secreted leptin from 3T3-L1 adipocytes treated with non-targeting (NT) or Exoc1 siRNA over 21 h. (**F**) Immunoblot showing that downregulation of *EXOC3* in a population of human JK6L lymphocytes correlates with a significant decrease in IgG secretion over a period of 24h. (**G**) Quantification of secreted heavy chain IgG over four independent experiments described in F (**H**) Quantification of secreted light chain IgG over four independent experiments described in F **P≤0.01; ***P≤0.001.

### Exocyst is essential for secretion in professional secretory cells

To test if exocyst is essential for constitutive secretion in other mammalian cells we tested this in highly specialised secretory cell models. Adipocytes have a well characterised regulated exocytosis pathway that is exocyst-dependent^45^, as well as a constitutive secretory pathway that is essential for the secretion of leptin and adiponectin that have also been associated with exocyst^46^. To test if secretion of these hormones was dependent on exocyst we abrogated expression of exocyst subunit *Exoc1* using siRNA in 3T3-L1 mouse adipocytes (Fig. 6C). We assayed the culture media using ELISA and observed a significant decrease in both adiponectin and leptin secretion (Fig. 6D).

Antibodies are constitutively secreted by blood lymphocytes. To test if exocyst is necessary for the secretion of antibodies in these cells, we used transient CRISPR-Cas9 to KO *EXOC3* in a clonal myeloma cell line that secretes complete IgG (both heavy and light antibody chains). We incubated the cells for 24 hours in fresh media and observed a significant decrease in antibody secretion by immunoblot (Fig. 6E-F). Together, these data demonstrate exocyst is essential for constitutive protein secretion across multiple different secretory cell types in mammalian cells.

## Discussion

In this study, we have identified the exocyst complex as an essential component of the mammalian secretory pathway. Exocyst localises to post-Golgi carriers and loss of exocyst prevents delivery of cell surface carriers causing them to accumulate in cell tips resulting in a global loss of secretion. In addition, in professional secretory cells such as adipocytes and lymphoma cells secretion of key proteins such as adipokines and antibodies is heavily reduced upon loss of exocyst.

The role of exocyst as the secretory complex in *Saccharomyces cerevisiae* is well established, due to its original identification in the *Sec* genetic screen^39^. Exocyst has been previously implicated in biosynthetic sorting in mammalian cells using *ts*VSV-G^27^ which colocalises with exocyst in the Golgi stacks, however, exocyst inhibition with antibodies did not affect VSV-G delivery^23,27^. siRNA depletion of *EXOC7* decreased the efficiency of *ts*VSV-G delivery to the plasma membrane^30^. Studies into post-Golgi carriers and thus secretion have been hampered by the lack of model systems to study the kinetic process. The use of quantitative RUSH assays in this study allows this route to be studied with kinetics that better resemble endogenous trafficking. In addition, loss of the exocyst complex is lethal in cultured cells^40^. To study knock-out of exocyst we have used transient CRISPR-Cas9. This has two key advantages, it allows abrogation of the protein-of-interest to be studied in the appropriate phenotypic window and it avoids artefacts introduced by clonal selection after CRISPR-Cas9 gene editing. The combination of kinetic trafficking assays and transient CRISPR-Cas9 thus provides new insights into the fundamental process of secretion.

Using the RUSH system coupled with kinetic trafficking assays, super-resolution imaging and TIRF microscopy we are able to map the machinery of the secretory pathway from the Golgi apparatus to the plasma membrane. RAB6 is essential for budding of the carriers from the Golgi (Fig. 2), where they traffic along microtubules to the cell tips. After budding, we see the tubules acquiring ARHGEF10 and RAB8 through imaging (Fig. 2), which is suggestive of a Rab cascade as previously described^24,35^. A RAB6 to RAB8 transition has also been described in the tethering and fusion of carriers via ELKS^24^. Across different species, both RAB8 and RAB11 have been described as EXOC6 interactors^47–50^. It has also been suggested that yeast homologues of RAB8 and RAB11 together with the exocyst complex can promote vesicle transport along the cytoskeleton via EXOC6 interaction with myosin type V^51,52^. We observe that GTPases RALA and RALB are redundantly important for the successful fusion of these carriers (Fig. 4), which have been shown in literature to bind both EXOC2 and EXOC8^26^. Besides the interaction with Rabs and Rals, EXOC5 has been shown to bind GTP-ARF6^53^, which has been recently proposed to mediate the recruitment of a PIP5 kinase^41^. In fact, three PIP5 kinases appear to act redundantly to convert the post-Golgi phospholipids (Fig. 4) and allow the recruitment of the exocyst complex, which tethers the carrier to the plasma membrane for the final fusion event. In mammalian cells, the PIP5 kinases are likely recruited to post-golgi membranes by several GTPases that act redundantly with Arf6. Indeed, this study has started to uncover a complex cargo delivery system, where redundancy most likely marks every step.

ELKS has previously been identified as the molecular tether for secretory carriers^22^. Although we observed ELKS localising to ‘hot-spots’ on the plasma membrane, we did not see a phenotype of *ELKS*-ko on cell surface delivery of LAMP1Δ-RUSH (Fig. S1). This does not rule out the role of ELKS in this process and it is possible that for certain cargoes or cell types ELKS has an essential role. Accordingly, we observe an increase in the phenotype when combining *ELKS* and *EXOC1* knock-out indicating a potential functional redundancy. Indeed, in neurons, studies demonstrate that ELKS acts as a redundant scaffold protein at the active zone site and that when this structure is disrupted, synaptic vesicle fusion is impaired^54^. Besides a structural role, ELKS has been further shown to capture RAB6 positive cargoes in golgin-like manner contributing to the establishment of a ready to fuse pool of synaptic vesicles at the active zone^25^. To fully understand the specific balance of roles between exocyst and ELKS will require further studies of other cargoes in specific cell types.

A number of exocyst subunits have been implicated in rare human genetic disorders. To date and to our knowledge there are four disease-associated subunits, EXOC8^55^ (MIM: 615283), EXOC7^55^ (MIM:608163) and EXOC4^56^ (MIM: 608185) and EXOC2^57^ (MIM: 615329). Mutations in EXOC8 and EXOC2 are associated with neurodevelopmental disorders and mutations in EXOC4 have been associated with nephrotic syndrome. In cell models exocyst has shown to be essential, with single-cell lethality associated with stable knock-outs^40^. The severity and rarity of the disorders associated with loss of exocyst subunits are likely linked to both the variety of cellular functions associated with exocyst as well as the fundamental nature of these processes, including secretion.

Exocyst was initially associated with basolateral vesicle trafficking to cell-cell contacts in polarised cells, and has since been implicated in a variety of processes including ciliogenesis, cytokinesis and the fusion of vesicles derived from recycling endosomes^26^. Here, we demonstrate that in addition to these roles, exocyst is essential for the fusion of secretory carriers. The regulation of exocyst and the recruitment of exocyst to carriers is not fully understood. There are a plethora of associated proteins that potentially allow for differential recruitment of exocyst to various carriers. These include the RALs^41^ (Fig. 4), ARF6^41^, phospholipids^41^ (Fig. 4), CDC42^58^, RAB10^59^, RAB11^60^ and RAB8A^61^. In addition, *EXOC3* has three homologues *EXOC3L1, EXOC3L2* and *EXOC3L4*; and *EXOC6* has *EXOC6B*. Which combination of these homologues or associated proteins provide specificity, redundancy or regulation of exocyst is not fully understood, but could potentially explain the widespread function of the complex with discrete specificities.

## Methods

### Antibodies and Other Reagents

The following primary antibodies were used for WB in this study: mouse anti-GFP HRP [GG4-2C2.12.10] (1:5000; 130-091-833; Miltenyi Biotec), rabbit anti-EXOC1 antibody (1:5000; ab118798; Abcam), rabbit anti-ELKS antibody [EPR13777] (1:5000; ab180507; Abcam), rabbit anti-TIMP2 antibody (D18B7) (1:1000; 5738; Cell Signaling Technology), rabbit anti-Cystatin C antibody [EPR4413] (1:5000; ab109508; Abcam), rabbit anti-PSAP antibody (1:1000; HPA004426; Atlas), mouse anti-GM130 antibody (1:1000; 610822; BD Transduction Laboratories), and rabbit anti-GAPDH HRP conjugate antibody (D16H11) (1:1000; 8884; Cell Signaling Technology). Goat HRP-conjugated secondary antibodies (1:5000) were purchased from Abcam (anti-mouse: ab205719; antirabbit: ab205718), and goat-anti-human IgG secondary antibody IRDye^®^ 800CW (1:15000; 926-32232) from LI-COR. The prokaryote expression vector encoding an anti-GFP mCherry nanobody (a gift from Martin Spiess, Addgene plasmid #109421) and pOPINE GFP nanobody:HALO:His6, encoding anti-GFP HALO nanobody (a gift from Lennart Wirthmueller, Addgene plasmid #111090) were expressed in bacteria and respectively GST- and His-purified in-house. The following cell-permeable dyes were obtained from these vendors: 646 HALO Dye (GA112A; Promega) and DAPI (D21490 Invitrogen).

### Plasmids

SBP-GFP-LAMP1ΔYQTI was PCR amplified from the original backbone SBP-GFP-LAMP11 (a generous gift from Juan Bonifacino, NICHD, NIH, USA) and Gibson assembled (E2621L; New England Biolabs) to the pEGFP-C1 (Clontech) vector backbone in between the AgeI/HindIII restriction sites. The SBP-LAMP1ΔYQTI-GFP was then obtained by a two step assembly that first removed GFP and then re-cloned its PCR product downstream of LAMP1ΔYQTI. Strep-KDEL was PCR amplified from Strep-KDEL-SBP-mCherry-GPI (Addgene plasmid #65295) and assembled to a BamHI/PsrI digested TtTMPV-Neo viral backbone (Addgene plasmid #27993). The neomycin resistance gene was then replaced with a hygromycin B encoding sequence.

pHALO-C1 and pHALO-N1 were generated by replacing the eGFP in Clontech vectors with HALO using Gibson assembly (E2621L; New England Biolabs). HALO-ARHGEF10, EXOC1-HALO, EXOC3-HALO, EXOC5-HALO, EXOC6-HALO, EXOC7-HALO, HALO-ELKS and HALO-EXOC6 were cloned by Gibson assembly using a synthetic, codon optimised version of each gene of interest (Integrated DNA Technologies), cloned in-frame upstream or downstream of HaloTag in pHALO-C1 or pHALO-N1. HALO-RAB6A and HALO-RAB8A were cloned in similar manner, except the gene sequences were PCR amplified from pEGFP-RAB6A (a kind gift from Juan Bonifacino, NICHD, NIH, USA) and pEGFP-RAB8A^62^ (a gift from Mitsunori Fukuda, Tohoku University, Japan) respectively. Point mutations were introduced via Q5 Site-Directed Mutagenesis Kit (M0554S; New England Biolabs).

Microtubules were visualised by expressing a plasmid containing β-tubulin-mCherry (Addgene plasmid #175829).

To generate a stable KO cell line, guide RNAs targeting ELKS (ERC1) were cloned into pSpCas9(BB)V2.0 (Addgene plasmid #62988)^63^, using the BbsI restriction sites.

For transient KO cells, guide RNAs targeting a gene of interest (Table S1) were cloned into pKLV-U6gRNA(BbsI)-PGKpuro2ABFP (Addgene plasmid #50946), using the BbsI restriction sites, as described above. The IDT Alt-R® CRISPR-Cas9 guide RNA tool was used to custom design two guide sequences per gene of interest. Cas9 viral expression backbone was a kind gift from Paul Lehner as well as the packaging vectors pMD.G and pCMVR8.91.

To generate the piggyback-RUSH constructs, we first created a piggyback-CMV-StrepKDEL-IRES-SBP-HALO. To achieve this, we Gibson assembled a PCR product containing CMV-StrepKDEL-IRES-SBP (amplified from Addgene plasmid #65295) to the SalI/MluI digested piggyback backbone (a generous gift from Jonathon Nixon-Abell, CIMR, University of Cambridge) and then digested the result with BamHI to assemble in a HaloTag flanked by BamHI and XbaI restriction sites. Gibson assembly was used to further clone other proteins of interest into this piggyback-RUSH-HALO vector. A PCR product of ColX (from Addgene #110726) and a synthetic gene containing PAUF (Integrated DNA Technologies) were cloned into the XbaI site downstream of HALO. PCR products containing CAB45 and NUCB1 (generous gifts from Liz Miller) were cloned into the BsrGI/BamHI sites upstream of HALO. Plasmids and primers used in this work are available upon request. All constructs were sequenced to verify their integrity.

### Cell lines

HeLa cells were already available in the lab; 3T3-L1 fibroblasts, originally from Howard Green (Harvard Medical School, Boston, MA), were a gift from David James (University of Sydney, Australia); JK6L myeloma cells were a kind gift from Jonathan Keats (Translational Genomics Research Institute, USA) and Lenti-X™ 293T cells were obtained from Takara Bio (632180). JK6L cells were maintained in RPMI 1640 (21875034; Gibco) supplemented with 10% fetal bovine serum (FBS) (F7424; Sigma-Aldrich), 1% GlutaMAX™ (35050061; Gibco) and MycoZap™ Plus-CL (VZA-2012; Lonza). They were kept in a humidified 5% CO2 atmosphere at 37°C. All other cell lines were grown in DMEM high glucose (D6429; Sigma-Aldrich) supplemented with 10% fetal bovine serum (FBS) (F7424; Sigma-Aldrich) and MycoZap™ Plus-CL (VZA-2012; Lonza), and were kept at 37°C in a humidified 5% CO2 atmosphere (10% CO2 for 3T3-L1 fibroblasts).

For differentiation to adipocytes, confluent 3T3-L1 fibroblasts were treated with 220 *μ*m dexamethasone (D4902; Sigma-Aldrich), 100 ng/mL biotin (B4501; Sigma-Aldrich), 2 mg/ml insulin (I5500; Sigma-Aldrich) and 500 mM IBMX (I5879; Sigma-Aldrich) in complete DMEM. After 3 days, media was replaced with fresh complete DMEM supplemented with 2 mg/ml insulin only, and after another 3 days (day 6 of differentiation), the media was then replaced with complete DMEM alone and subsequently changed every 48 hrs. A Hela cell line, stably expressing the Strep-KDEL was generated by infection with retroviral particles containing the Strep-KDEL plasmid followed by hygromycin B selection and subsequent single-cell clonal isolation.

Both HeLa stable cell lines expressing either SBP-LAMP1 ΔYQTI-GFP or SBP-GFP-LAMP1ΔYQTI were created by transient transfection of corresponding vector backbones using LipofectamineTM 2000 (11668019; Invitrogen), followed by selection with geneticin and single clonal isolation using live cell sorting. For SBP-LAMP1ΔYQTI-GFP two cell clones were selected: one low expression suitable for biochemical studies (i.e. CarrierIP) and another with higher expression to image. As for the SBP-GFP-LAMP1 ΔYQTI cell line, a medium to low expression clone was selected, suitable for the FACS LAMP1 cell surface assay. Lenti-XTM 293T cells were used to package the pKLV-puro vectors encoding plasmid/guide RNAs into lentiviral particles as previously described^64^. Viral supernatants were harvested after 48h, filtered through a 0.45 *μ*m filter, and when needed, concentrated down 10 times using the Lenti-X Concentrator (631232; Takara Bio). Supernatants were kept at −80°C prior to being directly applied to target cells which were subsequently spun at 700 xg for 1 h at 37°C. When necessary, cells were transiently selected with the appropriate antibiotic 48 h post-transduction.

Stable Cas9 expressing cell lines were generated by infecting target cells with lentiviral particles carrying Cas9 plasmid DNA (generous gift from Paul Lehner) followed by selection for blasticidin antibiotic resistance. Cas9 expression on >96% of the cell population was further confirmed through FACS analysis, by testing loss of cell surface expression of beta-2 microglobulin, upon transduction with lentiviral particles containing a B2M targeting sgRNA, using a mouse monoclonal anti-beta-2 microglobulin antibody (a generous gift from Paul Lehner).

### RUSH and FACS

10^6^ LAMP1Δ-RUSH HeLa cells were resuspended in DMEM supplemented with 25 mM HEPES (25 mM; 15630080; Gibco) and transferred to 1.5 ml microcentrifuge tubes. Tubes were kept at 37°C in a heating block (DB200/2; Techne) and D-biotin (B4501; Sigma-Aldrich) at a final concentration of 500 *μ*m was added at each time point (for the majority of experiments 35 min). Cells were then incubated on ice for 5 min to stop RUSH and subsequently spun down (4°C, 500 g, 5 min) to remove the supernatant. All downstream manipulations were either on ice or at 4°C. Following this, cells were incubated with an mCherry anti-GFP nanobody (10 ug/ml in PBS) for 1h, and then washed two times with 500 *μ*m of PBS. Finally, cells were filtered using the Cell-Strainer capped tubes (352235; FALCON). A minimum of 30.000 cells per sample was analysed using an LSRFortessa cell analyzer (BD Biosciences), gating for GFP-positive cells (indicative of LAMP1Δ-RUSH expression), plus any other concomitant fluorophore when appropriate. Data were analysed using FlowJo software (v10.8.1). RUSH was inferred by single-cell plotting the relative intensity of mCherry (LAMP1 at the PM) over that of GFP (LAMP1 available to RUSH).

### CRISPR knock-out

Stable ELKS1 KO HeLa cells were generated using the CRISPR-Cas9 system^63^. Plasmids containing Cas9 and single-guide RNAs were transfected (Lipofectamine 2000; 11668019; Invitrogen) into the LAMP1Δ-RUSH cell line according to manufacturers’ instructions. After 48 h, cells were selected with puromycin for a week before being single-cell sorted to establish clonal cell lines. Candidate clones were validated by immunoblotting to confirm the loss of ELKS.

Transient CRISPR KO was achieved by transducing previously established stable Cas9 cell lines. These include HeLa-Cas9, LAMP1 ΔYQTI-GFP-Cas9, SBP-GFP-LAMP1ΔYQTI-Cas9 and JK6L-Cas9. Briefly, 25×10^5^ LAMP1Δ-RUSH cells were transduced with 200 *μ*l of lentiviral supernatant or 30 *μ*l of 10x concentrated lentiviral particles in a 48-well plate. 48 h post-transduction cells were replated in a 6-well plate and incubated for an extra 4 days. On day 6, cells were detached with trypsin, and each well split into two, for two RUSH time points (0 and 35 min), followed by FACS analysis. Depletion of target protein was verified by immunoblotting and/or qPCR analysis. For some experiments, infections were scaled up accordingly. To recover the cell secretion phenotype, day 5 transient KO cells were electroporated (Gene Pulser Xcell Total System; Bio-Rad) with a plasmid encoding a guide resistant form of the protein of interest. In brief, 1×10^6^ cells were electroporated with 0.5 μg of plasmid DNA in a final volume of 200*μ*l of Opti-MEM I (31985062; Gibco), in a 2 mm electroporation cuvette (Z706086; Sigma-Aldrich), using the manufacturer’ settings for HeLa cells, and plated in fresh complete DMEM containing 646 HALO Dye (20 nM; GA112A; Promega) immediately after. The following day, cells were lifted, RUSHed for 35 min and FACS analysed as described above.

For *EXOC3* KO, 400×10^5^ JK6L-Cas9 cells were transduced with 30 *μ*l of 10x concentrated lentiviral particles in the presence of 0.8 *μ*m/ml polybrene (TR-1003; Sigma-Aldrich) and a final volume of 200 *μ*l of complete DMEM. This was carried out in a 48-well plate, without any centrifugation. The next day cells were replated in fresh complete RPMI and moved to a 6-well plate where they were incubated until the last day of the experiment. On day 2, puromycin was added (5 μg/ml) to select transduced myeloma cells. On day 5, cells were counted and seeded in a 96-well plate at a density of 2×10^6^/ml in complete fresh RPMI. 24 h later both the media and cell fractions were collected for immunoblot analysis of secreted IgG.

### siRNA

siRNA was delivered to differentiated adipocytes (6-7 days post-differentiation) using the TransIT-X2® dynamic delivery system (MIR 6004; Mirus Bio). Opti-MEM I (31985062; Gibco) and TransIT-X2® were mixed together (ratio 25/1). siRNA (siGENOME SMARTpool #69940; Dharmacon Horizon) was added to the Opti-MEM/TransIT-X2® mix such that the final concentration per well was 100 nM, mixed gently by pipetting, and incubated at room temperature for 30 min. 450 *μ*l of cell suspension was added to the Opti-MEM/ TransIT-X2®/siRNA mix and mixed well. 570 *μ*l of the resulting mixture was seeded into one well of a 24-well plate. Cells were fed with fresh culture media 24 hrs post-transfection and incubated at 37°C, 10% CO2 for a further 72 hrs. 96 h after initial transfection, cells were transfected with siRNA for a double-hit knockdown. Transfection mix was prepared as described above, except media alone was added to the Opti-MEM/TransIT-X2® mix instead of cell suspension. 570 mL of transfection mix was added to each well of adhered cells. Culture media was replaced with fresh media 24 h and 72 h post-transfection. 96 h after the second transfection, the culture media was harvested for analysis and cells were lysed for immunoblotting. Leptin and adiponectin present in the culture media were detected using the MSD® mouse leptin kit (MesoScale Discovery (MSD), Rockville, MD, USA, K152BYC-2) and the MSD® mouse adiponectin kit (MesoScale Discovery (MSD) Rockville, MD, USA, K152BYC-2) respectively.

### Carrier-IP

#### Preparation of GFP nanobody magnetic dynabeads

Magne® HaloTag® Beads (G7281; Promega) were incubated with purified HIS-HALO-GFP nanobody protein in 20 mM Hepes pH 7.5, 150 mM NaCl buffer overnight at 4°C. The beads were washed in the same buffer and stored at 4°C in 50 mM Hepes pH 7.5, 0.15 M NaCl, 15% (v/v) glycerol, 0.05% NaN3 (50% slurry).

Carrier IP: 1.75 x 10^6^ LAMP1 ΔYQTI-GFP-Cas9 cells were seeded in a 10 cm plate (per condition). 48h later, the cells were transfected with HALO-C1, HALO-RAB6a, EXOC1-HALO, EXOC3-HALO, EXOC5-HALO, EXOC6-HALO and EXOC7-HALO plasmids (Lipofectamine 2000; 11668019; Invitrogen) according to manufacturers’ instructions. 4 hours post-transfection media was replaced with fresh complete DMEM containing 646 HALO Dye (20 nM; GA112A; Promega). The next day, cells were washed twice with 10 ml PBS and detached with 3 ml of trypsin. Cells were collected with 2 times 10 ml of DMEM, spun down (4°C, 500 xg, 10 min) and resuspended in 20 ml of DMEM supplemented with 25 mM Hepes (25 mM; 15630080; Gibco). Tubes were kept at 37°C in a water bath and D-biotin (B4501; Sigma-Aldrich) was added at a final concentration of 500 *μ*m. Cells were incubated for 35 min at 37°C and then placed on ice, 20 ml of ice cold PBS was added immediately to stop RUSH. Cells were spun down (4°C, 500 xg, 10 min) to remove the supernatant, washed with 10 ml PBS and spun down again. Each pellet was resuspended with 1 ml of cytosol buffer (25 mM HEPES, 125 mM potassium acetate, 5.4 mM glucose, 25 mM magnesium acetate, adjust to pH7, and add fresh 100uM EDTA and 1x protease inhibitors) and the cells were homogenised with 25 strokes on ice using a cell homogeniser (8 micron bead; Isobiotec). Cell homogenates were spun at 3000 xg for 10 min at 4° C to remove cell debris. Supernatants were collected and 15 *μ*l were saved in 15 *μ*l of Laemmli buffer for the final gel. The rest of the supernatants were incubated for 15 min at 4°C with 10 *μ*l of GFP-Trap® Agarose beads (gta-10; Chromotek) to remove free GFP and spun at 1200 xg for 1 min at 4°C. Supernatants were then incubated with 30 *μ*l of homemade GFP nanobody magnetic dynabeads for 15 min at 4°C to trap

LAMP1Δ-RUSH carriers. The magnetic beads were washed 2 times with 1 ml of cytosol buffer (one quick wash and another for 5 min at 4°C) and 50 *μ*l of 2x Laemmli buffer. Samples were boiled for 10 min at 95°C and 10 *μ*l of the lysates and 20 *μ*l of the IP samples were resolved on a gradient Tris-Glycine acrylamide gel. The fluorescence of the 646-HaloTag ligand was directly imaged in the ChemiDoc Imaging System (Bio-Rad). Lamp1-GFP was detected by immunoblotting GFP.

### Microscopy

For immunofluorescence microscopy, 40×10^3^ day 6 transduced cells were plated onto Matrigel-coated (1:100 in complete DMEM; 354277; Corning) glass coverslips (400-03-19; Academy) in the presence of puromycin. The next day, coverslips were fixed with cytoskeletal fixing buffer (300 mM NaCl, 10 mM EDTA, 10 mM Glucose, 10 mM MgCl2, 20 mM PIPES pH 6.8, 2% Sucrose, 4% PFA) for 15 min and DAPI stained (300 nM; D21490 Invitrogen) for 5 min. PBS was used to wash cells in-between all steps and coverslips were mounted in ProLongTM Gold (P36930; Life Technologies). Standard fluorescent images were obtained with an Axio Imager.Z2 microscope (Zeiss) equipped with an Orca Flash 4.0 camera (Hamamatsu), an HXP 120V light source and a 100x NA 1.4 Plan-Apochromat objective, all under the control of ZEN software (Zeiss).

For live cell imaging, 0.9×10^6^ SBP-LAMP1 ΔYQTI-GFP cells were plated onto matrigel-coated glass coverslips (CB00250RAC; Menzel-Gläser). When required, FuGENE® 6 was used to transfect constructs encoding proteins of interest the following day. 2 hours post-transfection media was replaced with fresh complete DMEM containing 646 HALO Dye (20 nM; GA112A; Promega). The next day, cells were imaged in an Elyra 7 with Lattice-SIM microscope (Zeiss) equipped with an environmental chamber (temperature controlled at 37°C, humidified 5% CO2 atmosphere), two PCO.edge sC-MOS version 4.2 (CL HS) cameras (PCO), solid state diode continuous wave lasers and a Zeiss alpha Plan-Apochromat 63x/1.46 Oil Corr M27 objective for TIRF imaging and a Zeiss Plan-Apochromat 63x/1.4 Oil DIC M27 used for lattice-SIM, all under the control of ZEN black software (Zeiss).

### Secretomics

160×10^3^ HeLa-Cas9 cells were transduced with 1 ml of 1x EXOC3 lentiviral supernatant (total volume 2 ml) in a 6-well plate. The following day, media was replaced with fresh complete DMEM. 2 days later cells were replated into a 15 cm dish along with complete DMEM supplemented with puromycin, for EXOC3 KO condition. At infection day 7, both control and selected EXOC3 KO cells were washed three times with 30 ml PBS Ca+Mg+ (one quick wash, another for 5 min at 37°C, and a third quick one; D866; Sigma-Aldrich), and incubated with 15 ml of FBS free DMEM for 6 hours. The media was then collected and cooled on ice, and the cells lifted, counted sample buffer was added for a final concentration of 5×10^6^ cells/ml in 1x Laemmli buffer. Media was spun at 500 xg for 5 min at 4°C to exclude floating cells. Supernatant was then centrifuged at 4000 xg for 15 min at 4°C to exclude other cellular debris. The remaining media was then concentrated down to 500 uL using a falcon tube sized 3 kDa amicon column (UFC900308; Merck) spun at 4000 xg and 4°C for approximately 1 h. Concentrated media was analysed on a Q Exactive Orbitrap Mass Spectrometer (Thermo Scientific), and protein hits were filtered using a custom R script to select for proteins with a signal peptide, without a transmembrane domain and without a GPI anchor as per Uniprot annotations.

### Immunoblotting

Total denatured cell extracts (100.000 cells) were resolved on a gradient Tris-Glycine acrylamide gel and transferred to a PVDF membrane which was blocked with 5% skimmed milk and incubated with a primary antibody overnight at 4°C, followed by a 1 hour incubation with a secondary antibody at room temperature. Membranes were washed with PBS-T inbetween steps, and finally, their immunoreactivity was visualised using Clarity (1705061; Bio-Rad) or WesternBright Sirius (K-12043-D10; Advansta) ECL substrate, in the Chemi-Doc Imaging System (Bio-Rad). Bands were quantified using Image Lab software, version 6.1 (Bio-Rad) and GAPDH was used as a loading control.

### RNA extraction and qPCR

Total RNA was isolated from cultured WT or KO cells (35mm dish), using the RNeasy Mini Kit (74104; Qiagen) according to the manufacturer’s instructions. RNA quantification was performed using NanoDrop One (Thermo Scientific). Strand cDNA was generated by priming 1 μg of total RNA with a oligo(dT)16/random hexamers mix, using the High-Capacity RNA-to-cDNA™ Kit (4387406; Applied Biosistems), following manufacturer’s instructions. cDNA templates were diluted 10-fold and 1 *μ*l was used with specific oligos spanning 2 exons (Table S3) along with the PowerUpTM SYBRTM Green Master Mix (A25741; Applied Biosistems) for the qPCR reaction. All reactions were performed in 3 technical replicates using the CFX96 Touch Real-Time PCR Detection System (Bio-Rad). Data was analysed according to the 2-ΔΔCT method65.

### Statistical Analysis

Statistical analysis was performed using GraphPad Prism v.5 or Python 3.7 and statistical significance was considered when P<0.05. Comparisons were made using Student’s t-test or one-way ANOVA followed by Tukey post-hoc test for multiple comparisons. Unless stated, all quantitative data are expressed as mean ± standard deviation of at least three independent experiments.

## Acknowledgements

We thank Paul Lehner, Dick van den Boomen and Liz Miller for molecular biology reagents and advice. This research was supported by the CIMR Flow Cytometry Core Facility and the CIMR Microscopy Committee. In particular, we wish to thank Matthew Gratian, Reiner Schulte and Gabriela Grondys-Kotarba for their advice and support in imaging, flow cytometry and cell sorting D.G. and D.S. are funded by a Sir Henry Dale Fellowship awarded to D.G. from the Wellcome Trust/Royal Society (Grant 210481). CP is funded by a Wellcome Trust Institutional Strategic Support Fund (ISSF) awarded to DG and an Isaac Newton Trust Research Grant. DJF and AS were supported by a Medical Research Council Career Development Award to DJF (*MR/S*007091/1) This work was funded by a Wellcome and UKRI grant (Grant 210481, MR/S007091/1). For the purpose of open access, the author has applied a Creative Commons Attribution (CC BY) licence to any Author Accepted Manuscript version arising. Leptin and adiponectin secretion studies were supported by the MRC MDU Mouse Biochemistry Laboratory [*MCUU*00014/5], Cambridge. We thank Steve Royle and the Royle Lab for the aviliblity of the 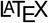 template used for typesetting. We thank Paul Luzio, Margaret Robinson and Laura Pellegrini for reading the manuscript, helpful discussions and advice.

## Supplementary Information

**Figure S1.**
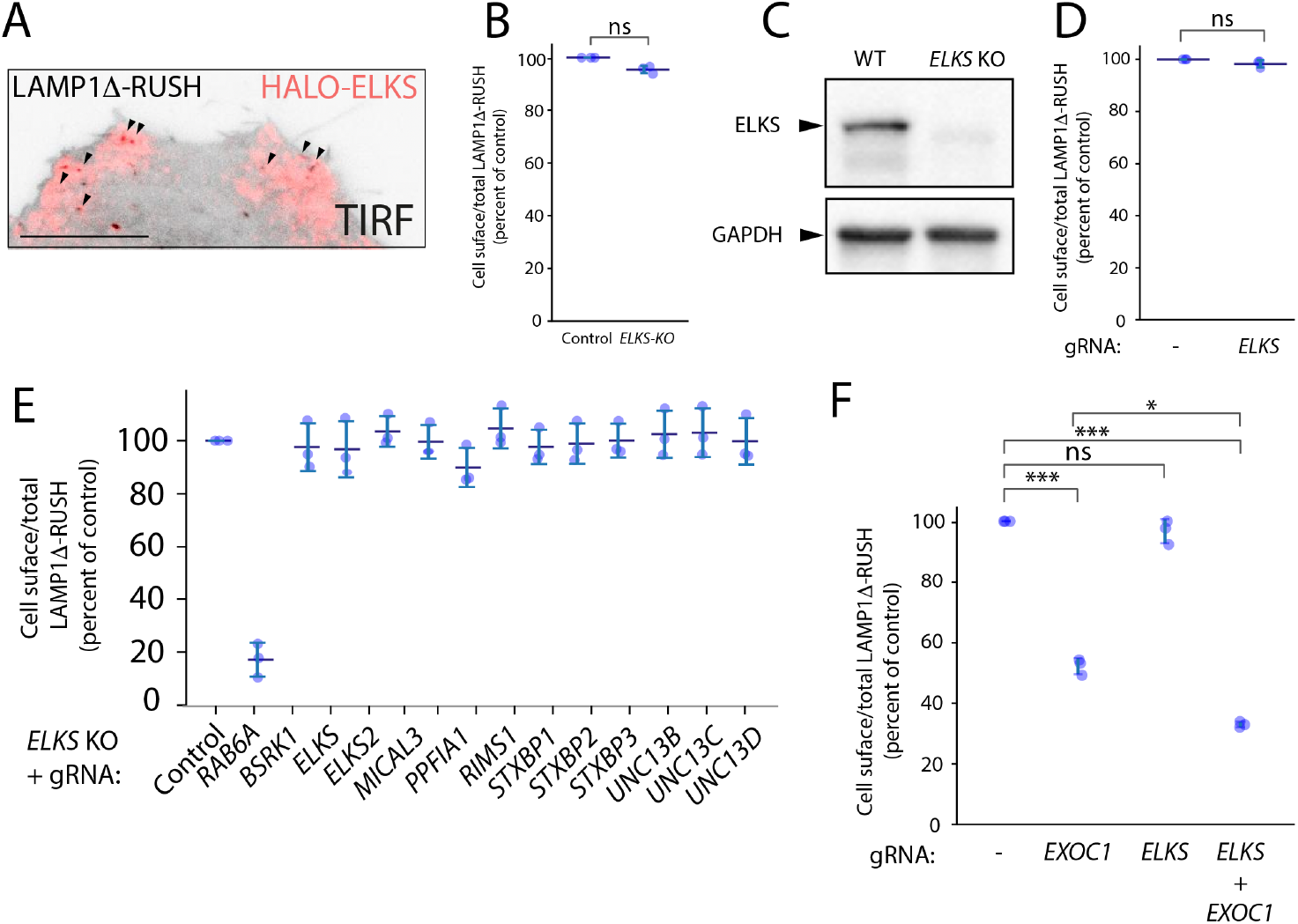
ELKS and its associated proteins and/or homologs are not necessary for LAMP1Δ-RUSH post-Golgi tubule fusion. (**A**) TIRF imaging of LAMP1Δ-RUSH (grey) cells expressing heterologous Halo-ELKS (red) mostly localised to fusion sites (arrowheads). Scale bar: 5 *μ*m. (**B**) Cell surface ratio quantification (FACS) of LAMP1Δ-RUSH at the plasma membrane after stable ELKS KO and 35 min of biotin exposure. (**C**) Immunoblot confirming loss of ELKS in a stable LAMP1Δ-RUSH ELKS KO clonal cell line. (**D**) Cell surface ratio quantification (FACS) of LAMP1Δ-RUSH at the plasma membrane after transient ELKS KO and 35 min of biotin exposure. (**E**) Cell surface ratio quantification (FACS) of LAMP1Δ-RUSH at the plasma membrane after transient candidate protein KO and 35 min of biotin exposure. This experiment was carried out in a stable clonal ELKS KO cell line.

**Figure S2.**
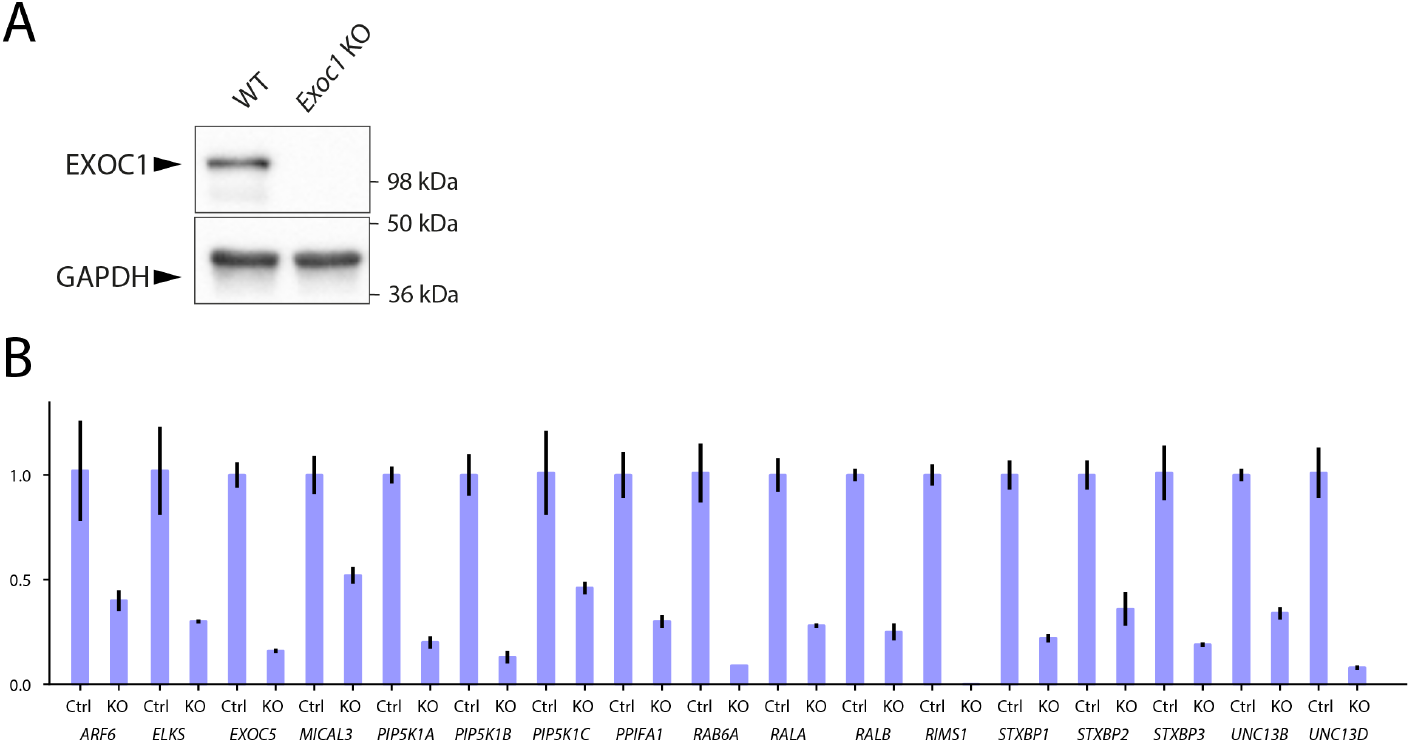
(**A**) Immunoblot confirming loss of EXOC1 in a transient LAMP1Δ-RUSH EXOC1 KO cell population. (**B**) Validation of guide RNA KO efficiency by qRT-PCR. Data from all conditions was internally normalised to GAPDH expression and is represented as fold change of control LAMP1Δ-RUSH Cas9 cells. ELKS2/ERC2 and UNC13C were not detectable by qPCR in control conditions.

**Figure S3.**
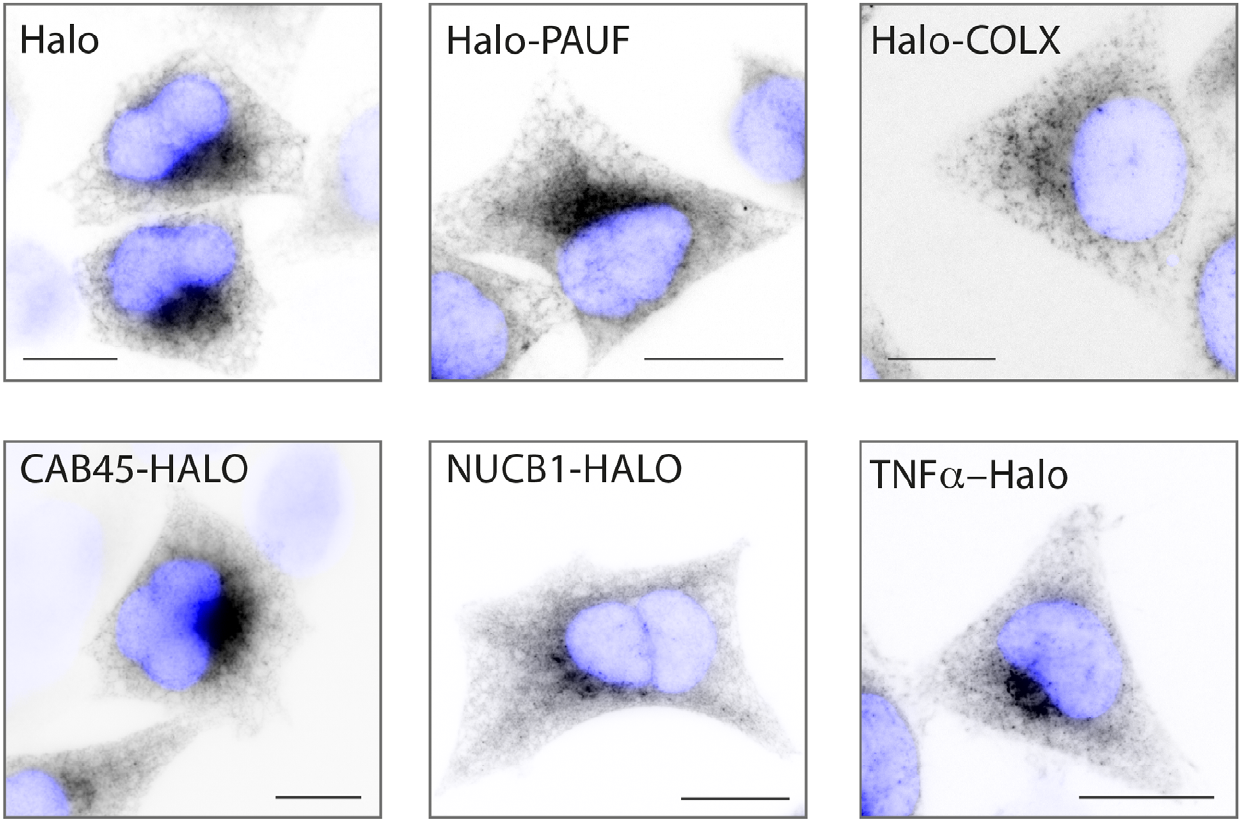
(**A**) Widefield imaging of stable cell lines expressing different RUSH cargos. Images show cargos effectively retained in the ER in the absence of biotin. Scale bar: 10 *μ*m.

## Supplementary Videos

**Figure SV1.**
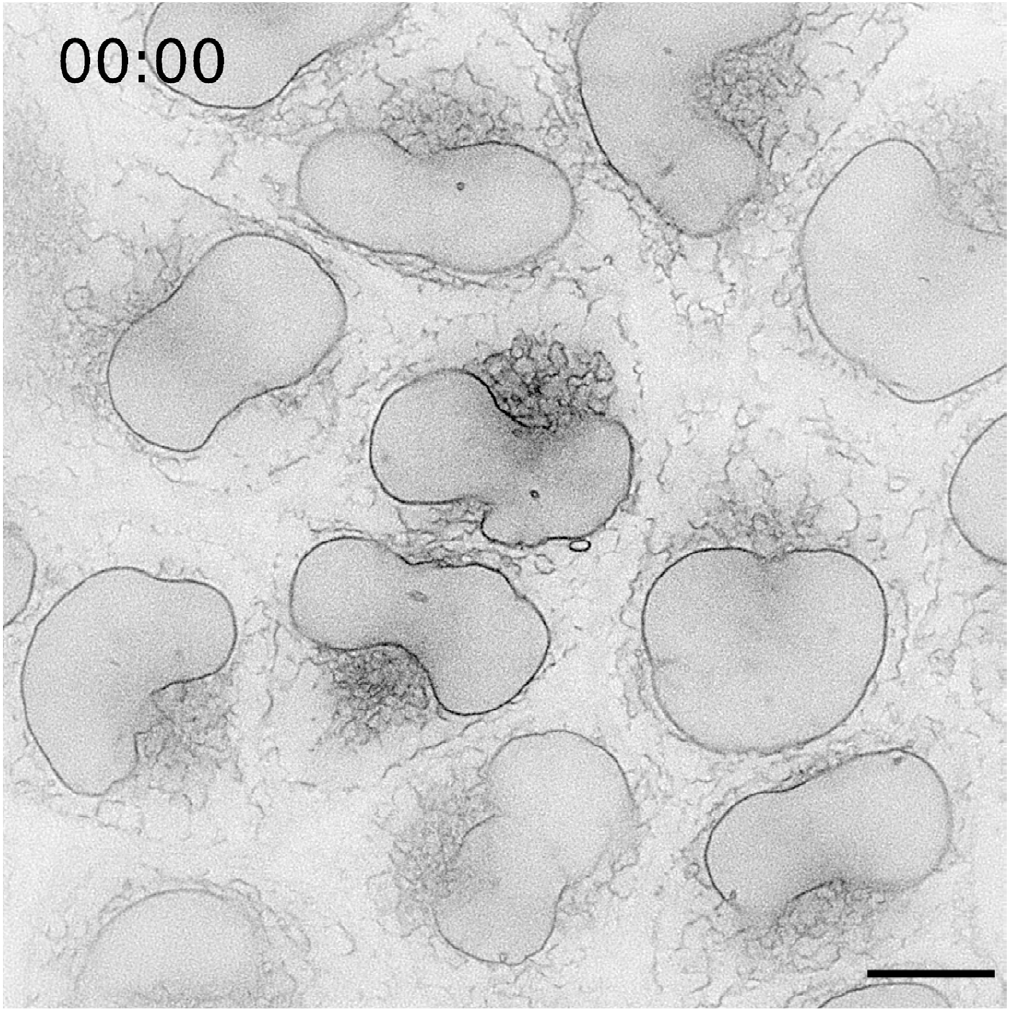
Supplementary Video 1B. Lattice-SIM live-cell imaging of LAMP1Δ-RUSH in HeLa cells. Cells were imaged every 30s for 1 hour after addition of biotin. Scale bar: 10 *μ*m

**Figure SV2.**
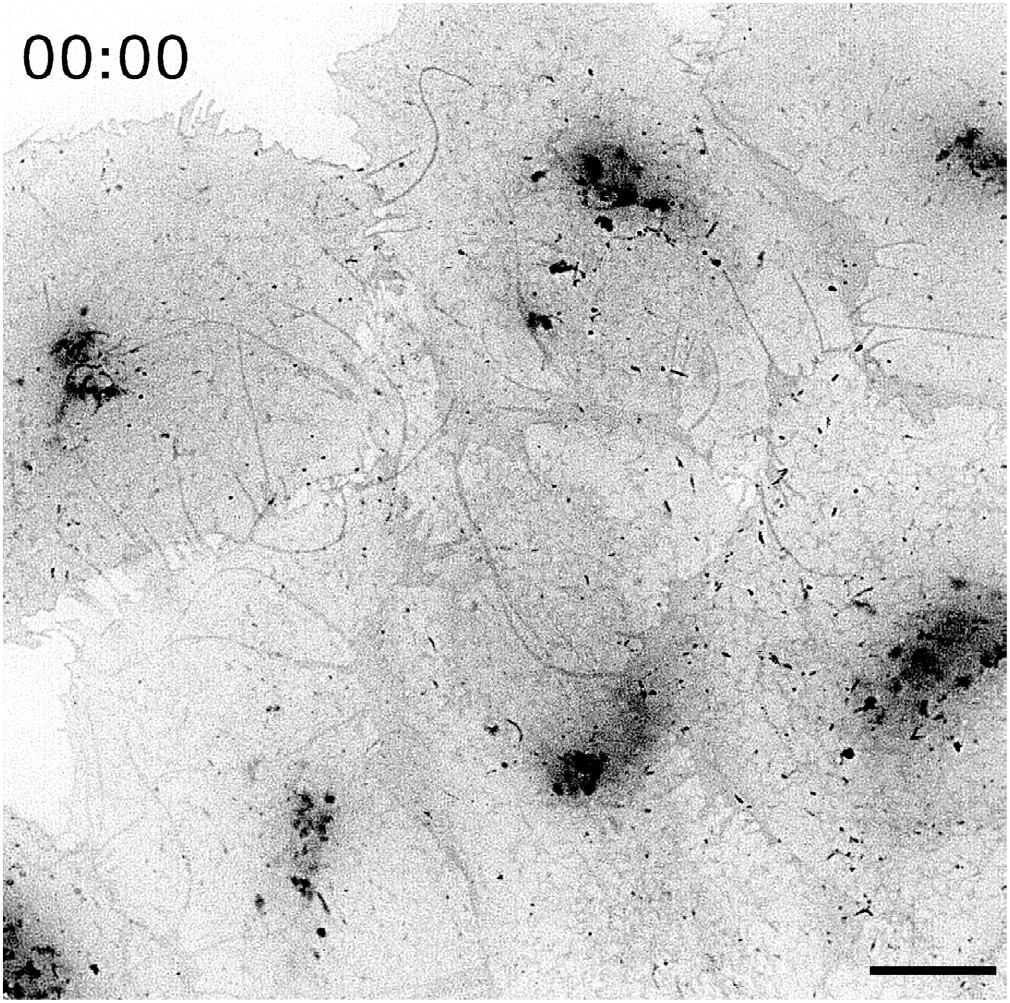
Supplementary Video 1C. Lattice-SIM live-cell imaging of LAMP1Δ-RUSH in HeLa cells 35 min after addition of biotin. Cells were imaged every 1.6 s for 5 min. Scale bar: 10 *μ*m.

**Figure SV3.**
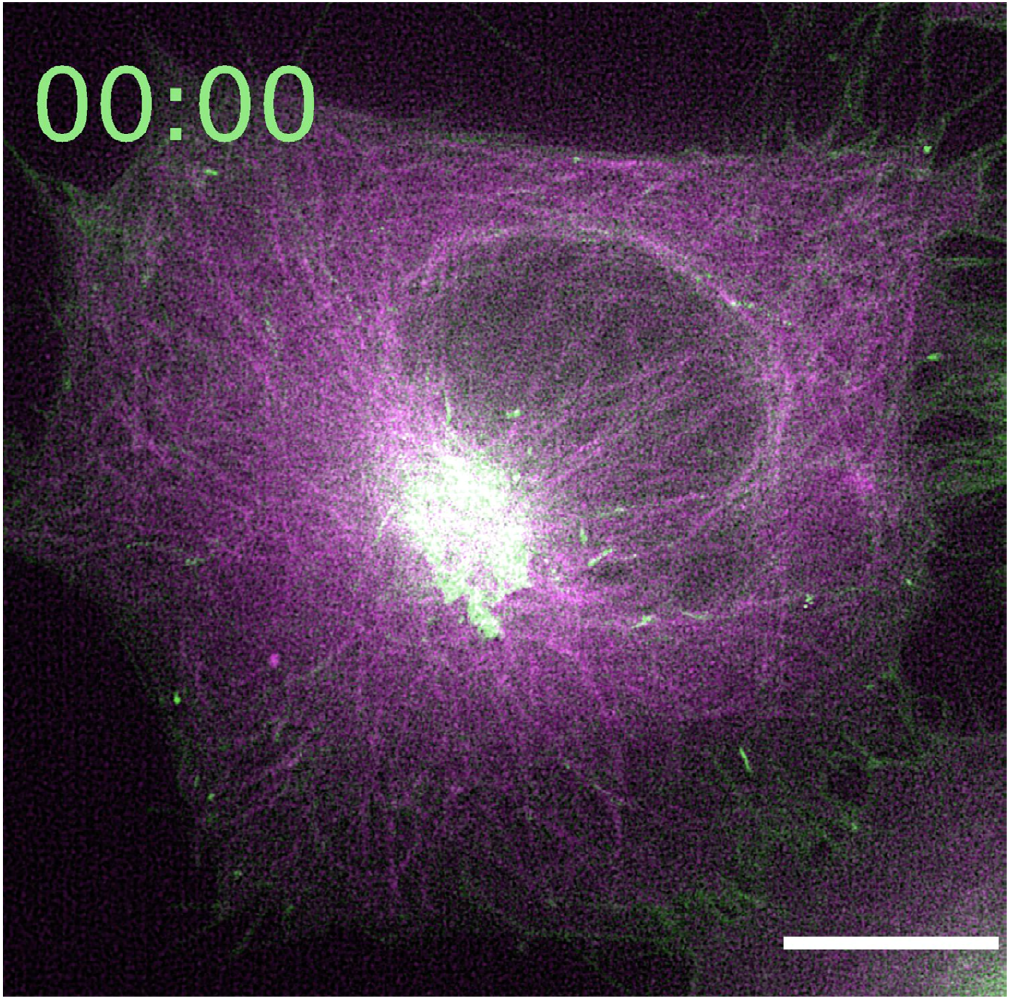
Supplementary Video 1D. Lattice-SIM live-cell imaging of a HeLa cell expressing stably LAMP1Δ-RUSH and transfected with β-tubulin-mCherry. The cell was imaged 34 min after addition of biotin every 1.6 s for 2.4 min. Scale bar: 10 *μ*m.

**Figure SV4.**
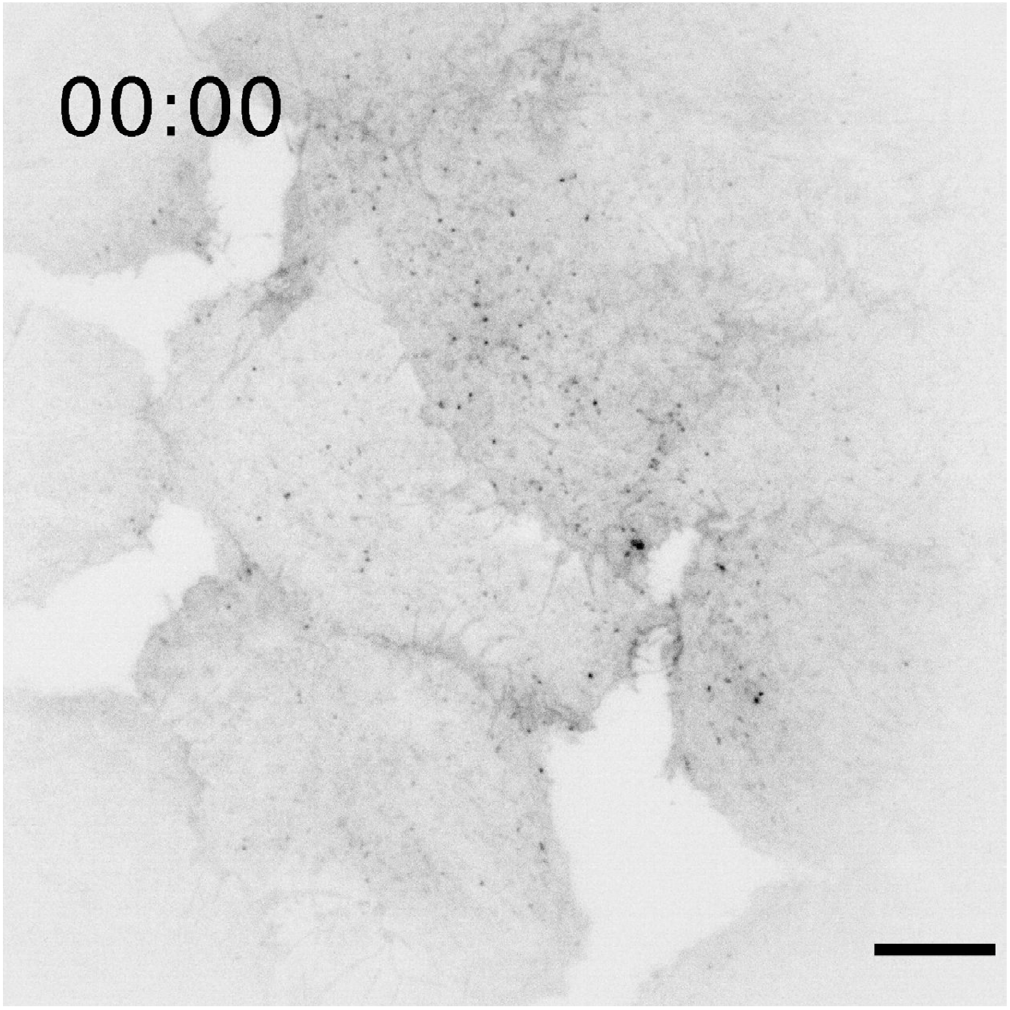
Supplementary Video 1E. ATIRF live-cell imaging of LAMP1Δ-RUSH in HeLa cells 35 min after addition of biotin. Cells were imaged every 30 ms for 7 s. Scale bar: 10 *μ*m

**Figure SV5.**
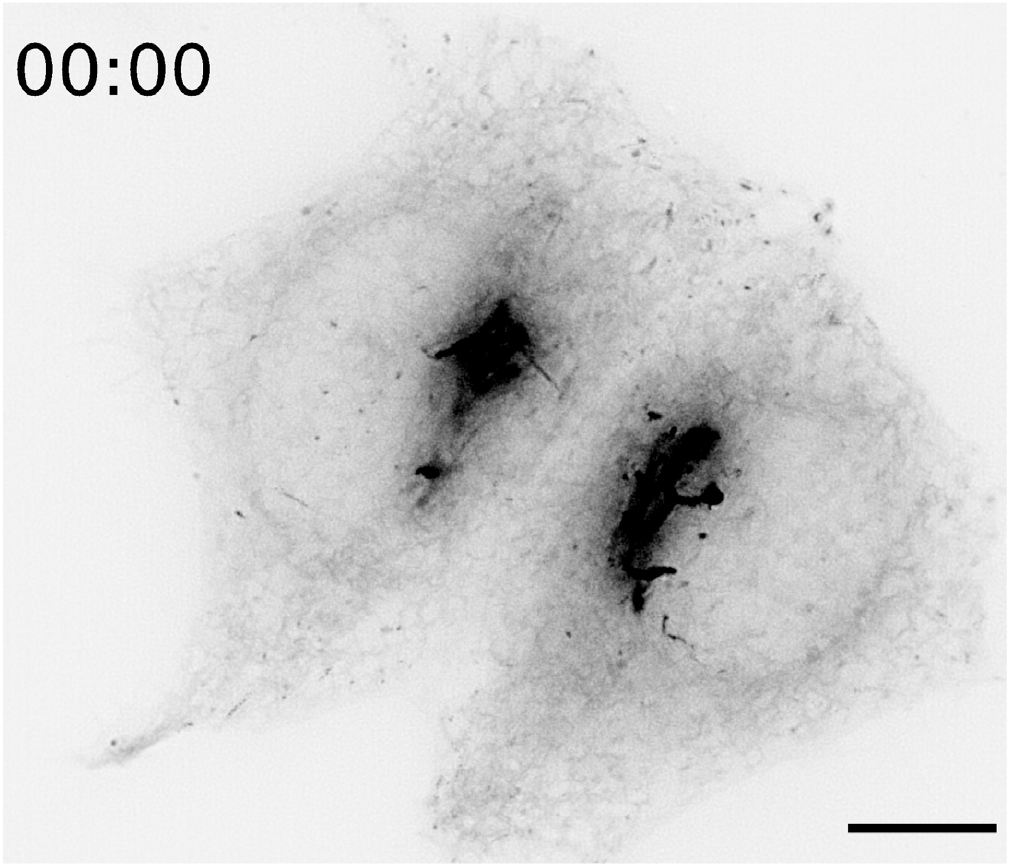
Supplementary Video 2A. Lattice-SIM live-cell imaging of HeLa cells overexpressing HALO-RAB6A. Cells were imaged every 1.6 s for 3.5 min. Scale bar: 10 *μ*m.

**Figure SV6.**
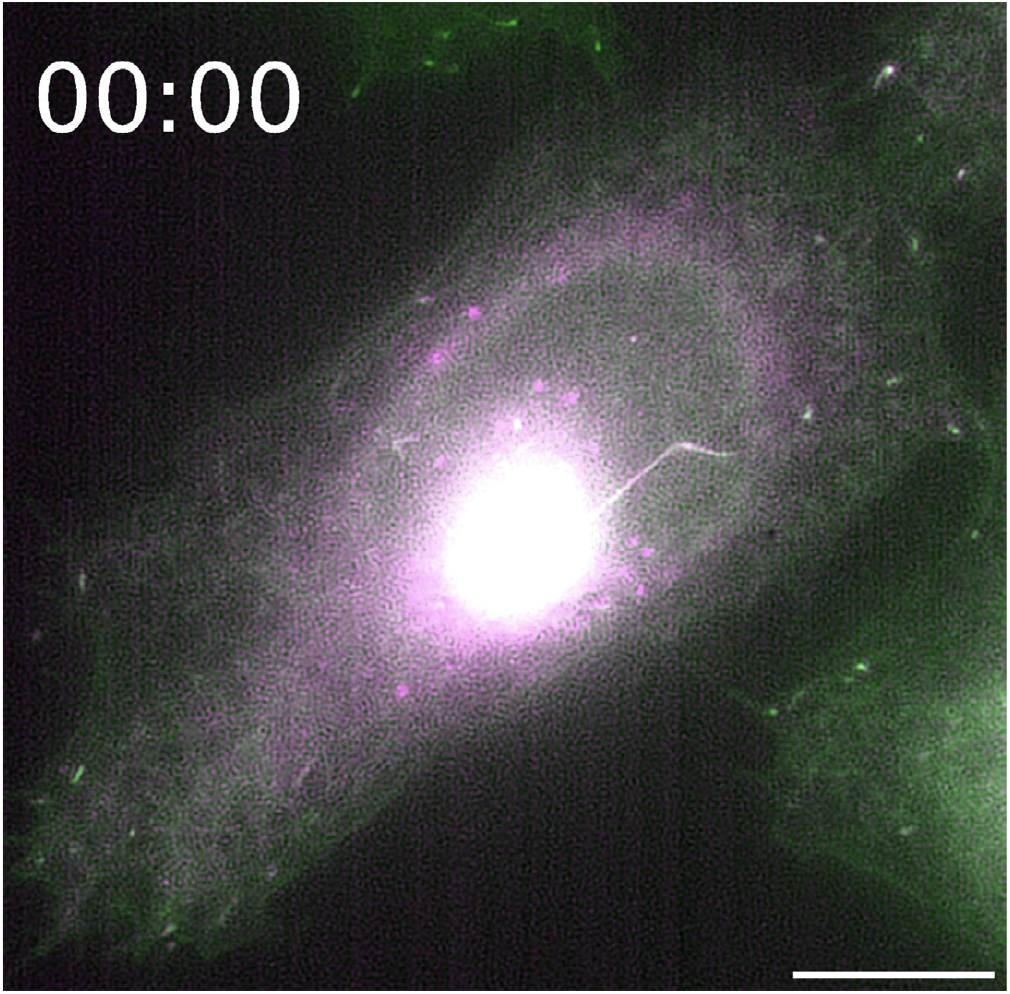
Supplementary Video 2B. Lattice-SIM live-cell imaging of a HeLa cell expressing stably LAMP1Δ-RUSH and transfected with HALO-RAB6A. The cell was imaged 21 min after addition of biotin every 3.16 s for 4.2 min. Scale bar: 10 *μ*m.

**Figure SV7.**
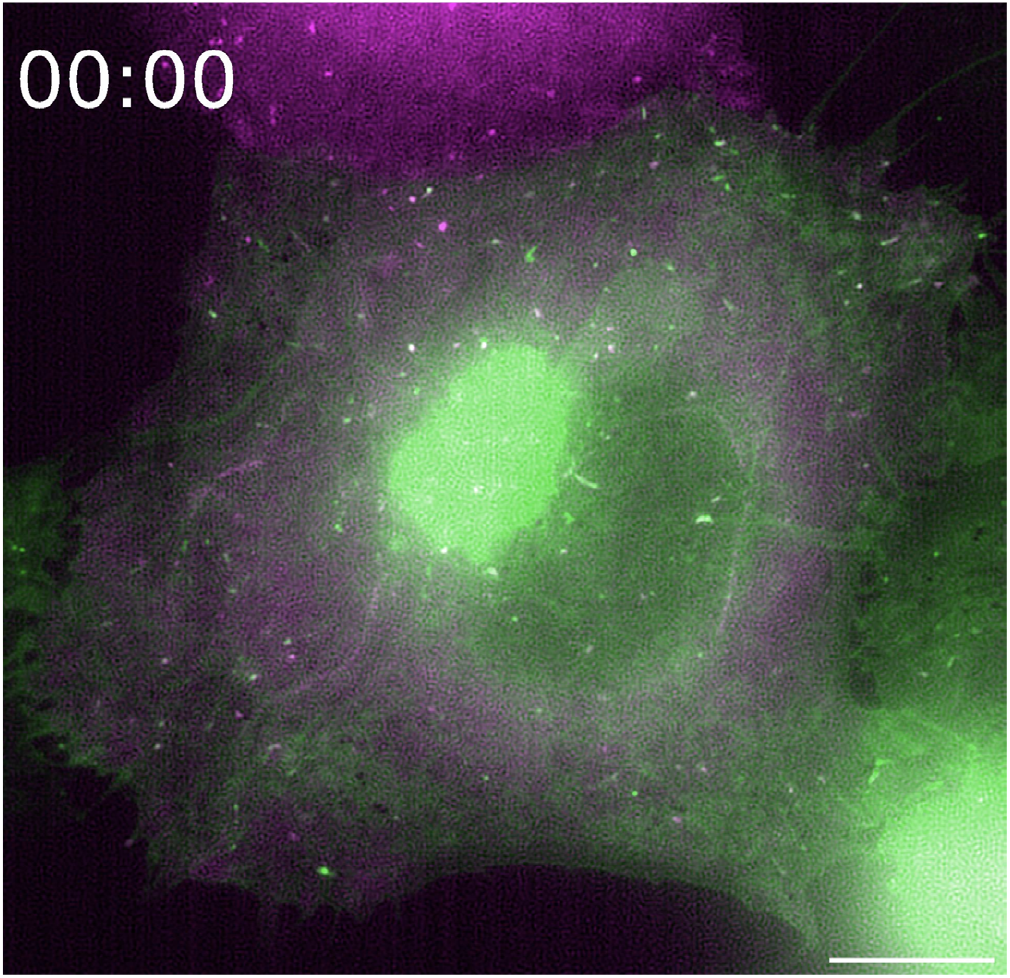
Supplementary Video 2C. Lattice-SIM live-cell imaging of a HeLa cell expressing stably LAMP1Δ-RUSH and transfected with HALO-ARHGEF10. The cell was imaged 32 min after addition of biotin every 3.16 s for 7.2 min. Scale bar: 10 *μ*m.

**Figure SV8.**
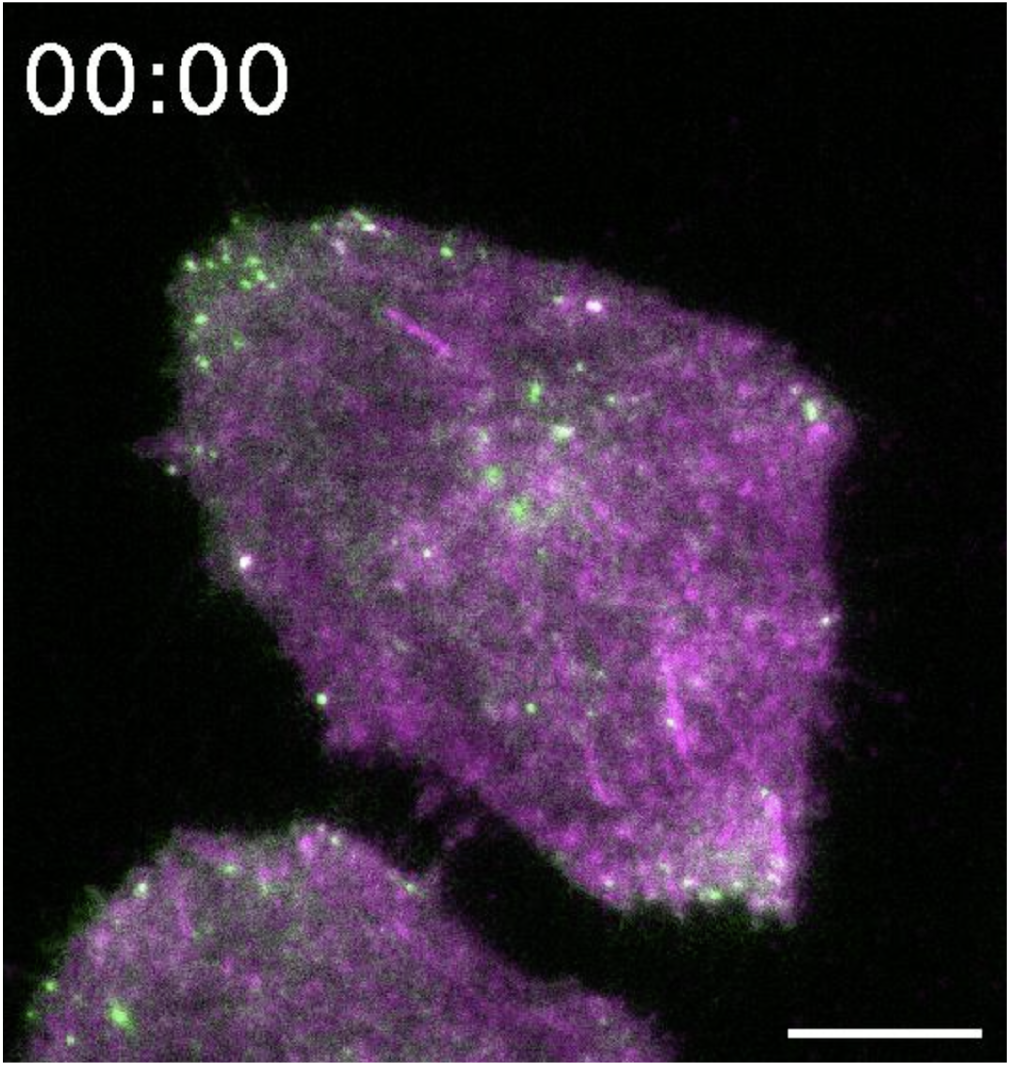
Supplementary Video 2D. A TIRF live-cell imaging of a HeLa cell expressing stably LAMP1Δ-RUSH and transfected with HALO-RAB8A. The cell was imaged 18 min after addition of biotin every 100 ms for 5.12 min. Scale bar: 10 *μ*m.

**Figure SV9.**
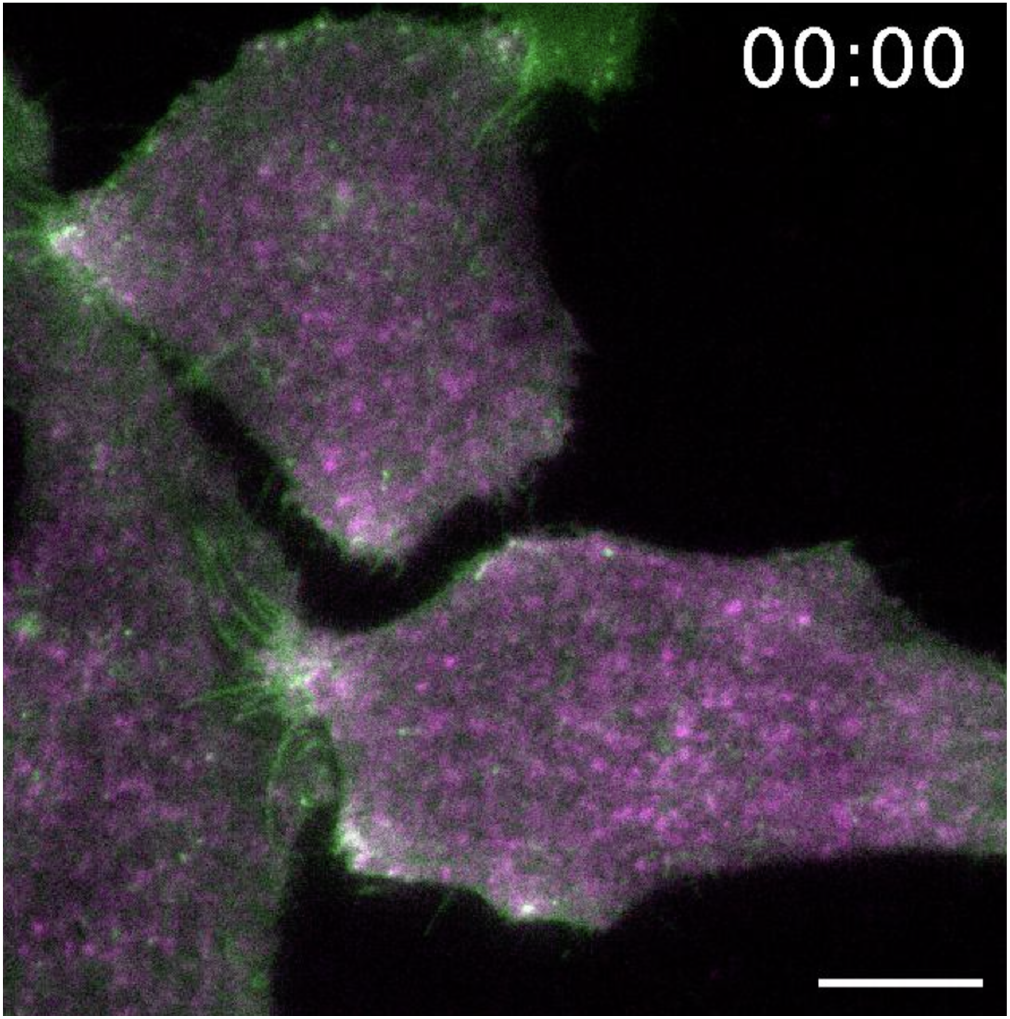
Supplementary Video 3A. TIRF live-cell imaging of a HeLa cell expressing stably LAMP1 Δ-RUSH and transfected with EXOC1-HALO. Cells were imaged 40 min after addition of biotin every 50 ms for 3.15 min. Scale bar: 10 *μ*m.

**Figure SV10.**
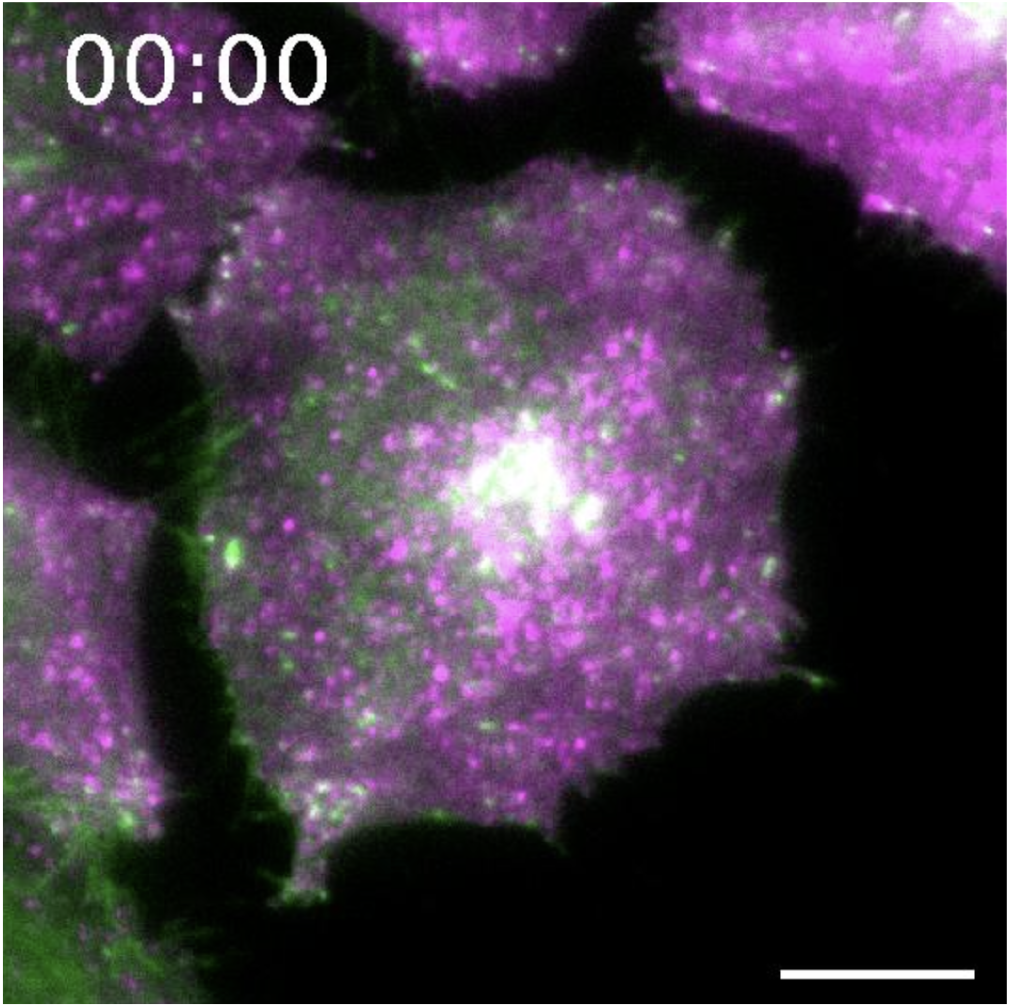
Supplementary Video 4A. TIRF live-cell imaging of a HeLa cell expressing stably LAMP1Δ-RUSH and transfected with HALO-EXOC6. Cells were imaged 17 min after addition of biotin every 110 ms for 6.2 min. Scale bar: 10 *μ*m.

**Figure SV11.**
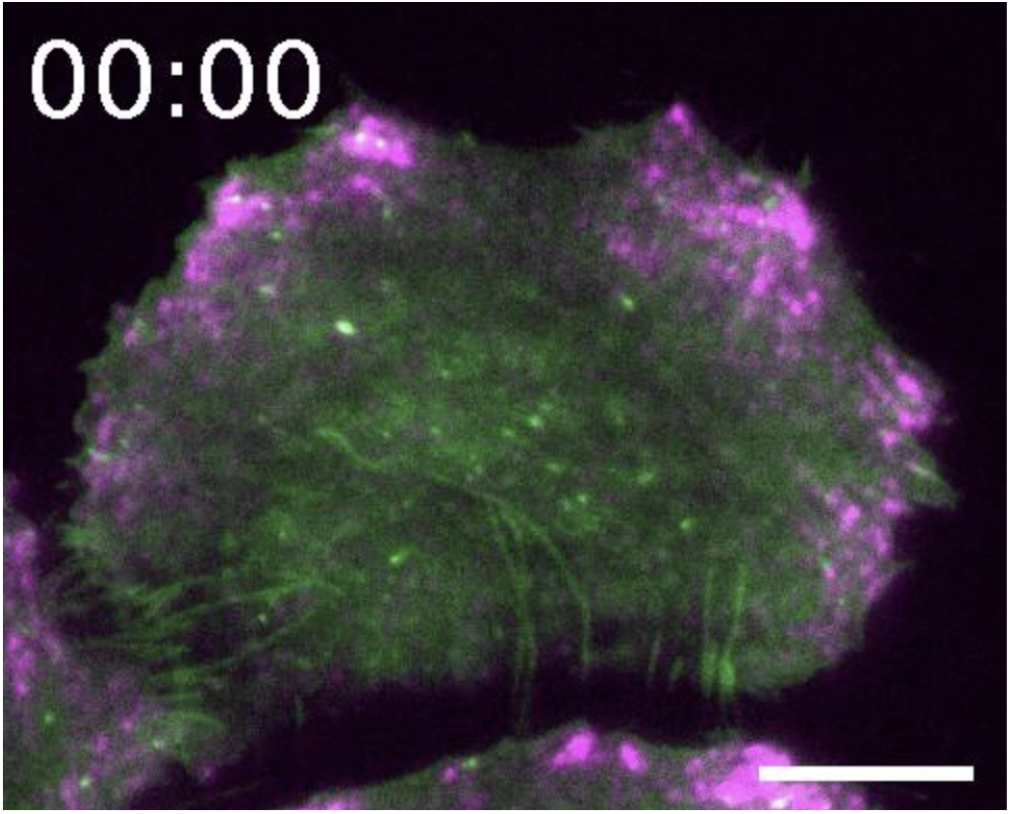
Supplementary Video S1A. TIRF live-cell imaging of a HeLa cell expressing stably LAMP1 Δ-RUSH and transfected with HALO-ELKS. Cells were imaged 31 min after addition of biotin every 110 ms for 3.4 min. Scale bar: 10 *μ*m.

## Bibliography

1. Chen, Y., Gershlick, D. C., Park, S. Y. & Bonifacino, J. S. Segregation in the Golgi complex precedes export of endolysosomal proteins in distinct transport carriers. J. Cell Biol. 216, 4141–4151 (2017).

2. Stalder, D. & Gershlick, D. C. Direct trafficking pathways from the Golgi apparatus to the plasma membrane. Semin. Cell Dev. Biol. 107, 112–125 (2020).

3. Kanapin, A. et al. Mouse proteome analysis. Genome Res. 13, 1335–1344 (2003).

4. Uhlén, M. et al. The human secretome. Sci. Signal. 12, (2019).

5. Thul, P. J. et al. A subcellular map of the human proteome. Science 356, (2017).

6. Xu, C. & Ng, D. T. W. Glycosylation-directed quality control of protein folding. Nat. Rev. Mol. Cell Biol. 16, 742–752 (2015).

7. Patterson, G. H. et al. Transport through the Golgi apparatus by rapid partitioning within a two-phase membrane system. Cell 133, 1055–1067 (2008).

8. Clermont, Y., Rambourg, A. & Hermo, L. Trans-Golgi network (TGN) of different cell types: three-dimensional structural characteristics and variability. Anat. Rec. 242, 289–301 (1995).

9. Keller, P., Toomre, D., Díaz, E., White, J. & Simons, K. Multicolour imaging of post-Golgi sorting and trafficking in live cells. Nat. Cell Biol. 3, 140–149 (2001).

10. Polishchuk, E. V., Di Pentima, A., Luini, A. & Polishchuk, R. S. Mechanism of constitutive export from the golgi: bulk flow via the formation, protrusion, and en bloc cleavage of large trans-golgi network tubular domains. Mol. Biol. Cell 14, 4470–4485 (2003).

11. Lupashin, V. & Sztul, E. Golgi tethering factors. Biochim. Biophys. Acta 1744, 325–339 (2005).

12. Whyte, J. R. & Munro, S. The Sec34/35 Golgi transport complex is related to the exocyst, defining a family of complexes involved in multiple steps of membrane traffic. Dev. Cell 1, 527–537 (2001).

13. Murray, D. H. et al. An endosomal tether undergoes an entropic collapse to bring vesicles together. Nature 537, 107–111 (2016).

14. Pérez-Victoria, F. J., Javier Pérez-Victoria, F. & Bonifacino, J. S. Dual Roles of the Mammalian GARP Complex in Tethering and SNARE Complex Assembly at the trans-Golgi Network. Molecular and Cellular Biology vol. 29 5251–5263 (2009).

15. Smith, R. D. & Lupashin, V. V. Role of the conserved oligomeric Golgi (COG) complex in protein glycosylation. Carbohydr. Res. 343, 2024–2031 (2008).

16. Spang, A. Membrane Tethering Complexes in the Endosomal System. Front Cell Dev Biol 4, 35 (2016).

17. Chou, H.-T., Dukovski, D., Chambers, M. G., Reinisch, K. M. & Walz, T. CATCHR, HOPS and CORVET tethering complexes share a similar architecture. Nature Publishing Group 23, 761–763 (2016).

18. Deguchi-Tawarada, M. et al. CAST2: identification and characterization of a protein structurally related to the presynaptic cytomatrix protein CAST. Genes Cells 9, 15–23 (2004).

19. Monier, S., Jollivet, F., Janoueix-Lerosey, I., Johannes, L. & Goud, B. Characterization of novel Rab6-interacting proteins involved in endosome-to-TGN transport. Traffic 3, 289–297 (2002).

20. Nakata, T. et al. Fusion of a novel gene, ELKS, to RET due to translocation t(10;12)(q11;p13) in a papillary thyroid carcinoma. Genes Chromosomes Cancer 25, 97–103 (1999).

21. Wang, Y., Liu, X., Biederer, T. & Südhof, T. C. A family of RIM-binding proteins regulated by alternative splicing: Implications for the genesis of synaptic active zones. Proc. Natl. Acad. Sci. U. S. A. 99, 14464–14469 (2002).

22. Fourriere, L. et al. RAB6 and microtubules restrict protein secretion to focal adhesions. J. Cell Biol. jcb.201805002 (2019).

23. Grigoriev, I. et al. Rab6 regulates transport and targeting of exocytotic carriers. Dev. Cell 13, 305–314 (2007).

24. Grigoriev, I. et al. Rab6, Rab8, and MICAL3 cooperate in controlling docking and fusion of exocytotic carriers. Curr. Biol. 21, 967–974 (2011).

25. Nyitrai, H., Wang, S. S. H. & Kaeser, P. S. ELKS1 Captures Rab6-Marked Vesicular Cargo in Presynaptic Nerve Terminals. Cell Rep. 31, 107712 (2020).

26. Wu, B. & Guo, W. The Exocyst at a Glance. J. Cell Sci. 128, 2957–2964 (2015).

27. Yeaman, C., Grindstaff, K. K., Wright, J. R. & Nelson, W. J. Sec6/8 complexes on trans-Golgi network and plasma membrane regulate late stages of exocytosis in mammalian cells. J. Cell Biol. 155, 593–604 (2001).

28. Ahmed, S. M. et al. Exocyst dynamics during vesicle tethering and fusion. Nat. Commun. 9, 5140 (2018).

29. Heider, M. R. & Munson, M. Exorcising the exocyst complex. Traffic 13, 898–907 (2012).

30. Liu, J., Zuo, X., Yue, P. & Guo, W. Phosphatidylinositol 4,5-bisphosphate mediates the targeting of the exocyst to the plasma membrane for exocytosis in mammalian cells. Mol. Biol. Cell 18, 4483–4492 (2007).

31. Boncompain, G. et al. Synchronization of secretory protein traffic in populations of cells. Nat. Methods 9, 493–498 (2012).

32. Munro, S. & Pelham, H. R. B. A C-terminal signal prevents secretion of luminal ER proteins. Cell 48, 899–907 (1987).

33. Buser, D. P., Schleicher, K. D., Prescianotto-Baschong, C. & Spiess, M. A versatile nanobody-based toolkit to analyze retrograde transport from the cell surface. Proc. Natl. Acad. Sci. U. S. A. 115, E6227–E6236 (2018).

34. Miserey-Lenkei, S. et al. Rab and actomyosin-dependent fission of transport vesicles at the Golgi complex. Nat. Cell Biol. 12, 645–654 (2010).

35. Shibata, S. et al. ARHGEF10 directs the localization of Rab8 to Rab6-positive executive vesicles. J. Cell Sci. 129, 3620–3634 (2016).

36. Bowser, R. & Novick, P. Sec15 protein, an essential component of the exocytotic apparatus, is associated with the plasma membrane and with a soluble 19.5S particle. J. Cell Biol. 112, 1117–1131 (1991).

37. Bowser, R., Müller, H., Govindan, B. & Novick, P. Sec8p and Sec15p are components of a plasma membrane-associated 19.5 S particle that may function downstream of Sec4p to control exocytosis. J. Cell Biol. 118, 1041–1056 (1992).

38. TerBush, D. R. & Novick, P. Sec6, Sec8, and Sec15 are components of a multisubunit complex which localizes to small bud tips in Saccharomyces cerevisiae. J. Cell Biol. 130, 299–312 (1995).

39. Novick, P. J., Field, C. & Schekman, R. Identification of 23 complementation groups required for post-translational events in the yeast secretory pathway. Cell 21, 205–215 (1980).

40. Wang, T. et al. Identification and characterization of essential genes in the human genome. Science 350, 1096–1101 (2015).

41. Maib, H. & Murray, D. H. A mechanism for exocyst-mediated tethering via Arf6 and PIP5K1C driven phosphoinositide conversion. bioRxiv 2021.10.14.464363 (2021) doi:10.1101/2021.10.14.464363.

42. Wakana, Y. et al. A new class of carriers that transport selective cargo from the trans Golgi network to the cell surface. EMBO J. 31, 3976–3990 (2012).

43. Pakdel, M. & von Blume, J. Exploring new routes for secretory protein export from the trans-Golgi network. Mol. Biol. Cell 29, 235–240 (2018).

44. Itzhak, D. N., Tyanova, S., Cox, J. & Borner, G. H. Global, quantitative and dynamic mapping of protein subcellular localization. Elife 5, (2016).

45. Inoue, M., Chang, L., Hwang, J., Chiang, S.-H. & Saltiel, A. R. The exocyst complex is required for targeting of Glut4 to the plasma membrane by insulin. Nature 422, 629–633 (2003).

46. Kuramoto, K., Kim, Y.-J., Hong, J. H. & He, C. The autophagy protein Becn1 improves insulin sensitivity by promoting adiponectin secretion via exocyst binding. Cell Rep. 35, 109184 (2021).

47. Zhang, X.-M., Ellis, S., Sriratana, A., Mitchell, C. A. & Rowe, T. Sec15 Is an Effector for the Rab11 GTPase in Mammalian Cells*. J. Biol. Chem. 279, 43027–43034 (2004).

48. Guo, W., Roth, D., Walch-Solimena, C. & Novick, P. The exocyst is an effector for Sec4p, targeting secretory vesicles to sites of exocytosis. EMBO J. 18, 1071–1080 (1999).

49. Wu, S., Mehta, S. Q., Pichaud, F., Bellen, H. J. & Quiocho, F. A. Sec15 interacts with Rab11 via a novel domain and affects Rab11 localization in vivo. Nat. Struct. Mol. Biol. 12, 879–885 (2005).

50. Luo, G., Zhang, J. & Guo, W. The role of Sec3p in secretory vesicle targeting and exocyst complex assembly. Mol. Biol. Cell 25, 3813–3822 (2014).

51. Lipatova, Z. et al. Direct interaction between a myosin V motor and the Rab GTPases Ypt31/32 is required for polarized secretion. Mol. Biol. Cell 19, 4177–4187 (2008).

52. Jin, Y. et al. Myosin V transports secretory vesicles via a Rab GTPase cascade and interaction with the exocyst complex. Dev. Cell 21, 1156–1170 (2011).

53. Prigent, M. et al. ARF6 controls post-endocytic recycling through its downstream exocyst complex effector. J. Cell Biol. 163, 1111–1121 (2003).

54. Wang, S. S. H. et al. Fusion Competent Synaptic Vesicles Persist upon Active Zone Disruption and Loss of Vesicle Docking. Neuron 91, 777–791 (2016).

55. Coulter, M. E. et al. Regulation of human cerebral cortical development by EXOC7 and EXOC8, components of the exocyst complex, and roles in neural progenitor cell proliferation and survival. Genet. Med. 22, 1040–1050 (2020).

56. Nihalani, D. et al. Disruption of the exocyst induces podocyte loss and dysfunction. J. Biol. Chem. 294, 10104–10119 (2019).

57. Van Bergen, N. J. et al. Mutations in the exocyst component EXOC2 cause severe defects in human brain development. J. Exp. Med. 217, (2020).

58. Zhang, X. et al. Cdc42 interacts with the exocyst and regulates polarized secretion. J. Biol. Chem. 276, 46745–46750 (2001).

59. Babbey, C. M., Bacallao, R. L. & Dunn, K. W. Rab10 associates with primary cilia and the exocyst complex in renal epithelial cells. Am. J. Physiol. Renal Physiol. 299, F495–506 (2010).

60. Takahashi, S. et al. Rab11 regulates exocytosis of recycling vesicles at the plasma membrane. J. Cell Sci. 125, 4049–4057 (2012).

61. Mei, K. & Guo, W. The exocyst complex. Curr. Biol. 28, R922–R925 (2018).

62. Matsui, T., Itoh, T. & Fukuda, M. Small GTPase Rab12 regulates constitutive degradation of transferrin receptor. Traffic 12, 1432–1443 (2011).

63. Ran, F. A. et al. Genome engineering using the CRISPR-Cas9 system. Nat. Protoc. 8, 2281–2308 (2013).

64. Pirona, A. C., Oktriani, R., Boettcher, M. & Hoheisel, J. D. Process for an efficient lentiviral cell transduction. Biol Methods Protoc 5, bpaa005 (2020).

65. Livak, K. J. & Schmittgen, T. D. Analysis of relative gene expression data using real-time quantitative PCR and the 2(-Delta Delta C(T)) Method. Methods 25, 402–408 (2001).

